# BiCLUM: Bilateral Contrastive Learning for Unpaired Single-Cell Multi-Omics Integration

**DOI:** 10.1101/2025.03.17.643621

**Authors:** Yin Guo, Izaskun Mallona, Mark D. Robinson, Limin Li

## Abstract

The integration of single-cell multi-omics data provides a powerful approach for understanding the complex interplay between different molecular modalities, such as RNA expression, chromatin accessibility and protein abundance, measured through assays like scRNA-seq, scATAC-seq and CITE-seq, at single-cell resolution. However, most existing single-cell technologies focus on individual modalities, limiting a comprehensive understanding of their interconnections. Integrating such diverse and often unpaired datasets remains a challenging task due to unknown cell correspondences across distinct feature spaces and limited insights into cell-type-specific activities in non-scRNA-seq modalities. In this work, we propose BiCLUM, a <u>Bi</u>lateral <u>C</u>ontrastive <u>L</u>earning approach for <u>U</u>npaired single-cell <u>M</u>ulti-omics integration, which simultaneously enforces cell-level and feature-level alignment across modalities. BiCLUM first transforms one modality, such as scATAC-seq, into the data space of another modality, such as scRNA-seq, using prior genomic knowledge. It then learns cell and gene embeddings simultaneously through a bilateral contrastive learning framework, incorporating both cell-level and feature-level contrastive losses. We evaluated BiCLUM on aligning gene expression with chromatin accessibility via three paired RNA-ATAC multi-omics datasets, as well as gene expression with protein expression via three CITE-seq datasets. The results demonstrate that BiCLUM either outperforms or is at least comparable to existing integration methods, excelling in both visualization and quantitative metrics. Furthermore, BiCLUM preserves the biological relevance of the integrated data, making it a potential powerful tool for downstream biological analysis, such as cell type identification and pathway exploration.

## Introduction

In recent years, single-cell analysis has emerged as a pivotal focus in biomedicine, driven by the development of advanced technologies designed to measure various molecular modalities at the single-cell level. For example, scRNA-seq has been widely adopted to profile gene expression [41, 35, 62], while chromatin accessibility [8, 18], DNA methylation [34, 37], and histone modifications [45] have also been explored at single-cell resolution. However, they often focus on individual modalities, limiting the full understanding of the relationships between them. To address this limitation, multimodal technologies [5, 43, 10] have been developed to enable the simultaneous measurement of multiple modalities within the same cells. By integrating these modalities, a more comprehensive understanding of the relationships across various cellular dimensions can be achieved, facilitating the construction of informative representations of cells.

The integration of single-cell multi-omics datasets can be broadly categorized into paired and unpaired scenarios.

In the paired scenario, multiple modalities are measured from the same cells. For example, Mowgli [26] is a joint integration approach that is based on integrative Nonnegative Matrix Factorization and Optimal Transport, specifically designed for such datasets. However, despite advancements in technologies capable of simultaneously profiling multiple modalities, conducting multi-omics experiments remains technically challenging and resource-intensive. As a result, most single-cell multi-omics datasets are generated through separate experiments involving different cell batches, leading to unpaired datasets. This poses significant challenges in integrating datasets across individuals, technologies, and species. The challenges of unpaired integration of multi-omics datasets primarily fall into two aspects. Firstly, different modalities lack common features, and the correspondences between features and cells are unknown. Secondly, compared to scRNA-seq data, the cell-type-specific activity of features in other modalities is less well established, posing additional challenges for effective integration.

Several methods have been developed to address the integration of single-cell multi-omics datasets in unpaired scenarios, which can be broadly classified into two categories. The first category of approaches focuses on aligning cells across different omics modalities using nonlinear manifold alignment techniques. Different methods in this category align the modalities either before embedding learning (e.g., UnionCom [11], Pamona [13]), after embedding learning (e.g., scTopoGAN [48]), or simultaneously during embedding learning (e.g., MMD- MA [49], JointMDS [16]). For example, UnionCom [11] aligns cells by matching geometric distance matrices of raw data across modalities, then projects them into a shared low-dimensional space where cells with inferred correspondences exhibit similar expression patterns. scTopoGAN [48] employs topological autoencoders to extract latent representations of each modality separately and utilizes a topology-guided generative adversarial network (GAN) to align these representations in a common feature space. JointMDS [16] integrates multidimensional scaling (MDS) with Wasserstein Procrustes analysis to simultaneously derive low-dimensional embeddings and infer correspondences across modalities. These integration methods for unpaired single-cell multi-omics datasets leverage a variety of innovative techniques combining latent embedding learning and manifold alignment to provide meaningful insights into cellular and molecular relationships. However, these manifold alignment-based methods may not fully consider biological relationships between modalities, potentially resulting in outcomes that lack biological interpretability.

The second category of approaches involves converting multi-omics datasets into a unified feature space, often leveraging prior biological knowledge. This common representation enables the integration of diverse datasets by highlighting shared patterns or features across modalities. Most existing methods in this category focus on integrating scRNA-seq and scATAC-seq datasets [32, 27, 61] by leveraging prior genomic information to convert ATAC features into RNA feature space. Specifically, these approaches convert scATAC-seq data into gene activity scores, thereby generating shared features (i.e., genes). For example, the Seurat3 method [52] identifies anchors using Canonical Correlation Analysis (CCA) and Mutual Nearest Neighbors (MNN), followed by filtering, scoring, and weighting these anchors to obtain a corrected expression matrix. The LIGER method [32] aligns datasets by identifying shared and dataset-specific factors through integrative non-negative matrix factorization. These two methods employ linear transformations, which may not sufficiently capture the complex topological structures inherent in different modalities. bindSC [20] adopts a different approach by using a transformed gene activity expression matrix as a bridge between multi-omics datasets and applying canonical correlation analysis (CCA) to align datasets at both the feature and cell levels. Some methods, instead of directly transforming features across modalities, leverage genomic information to establish connections between features. For example, GLUE [14] learns latent embeddings through a guidance graph to link features across modalities and uses an adversarial network to align cells. The guidance graph is a graph-based structure that leverages prior biological knowledge to establish relationships between features across different modalities, facilitating their integration. Similarly, scDART [61] uses prior knowledge to define a linkage matrix between features across modalities and aligns datasets via Maximum Mean Discrepancy (MMD). However, the method requires computing similarity matrices for each modality, which can cause memory issues with large datasets.

Another limitation is that most existing integration approaches are specifically designed for scRNA-seq and scATAC-seq datasets, thereby limiting their applicability to other modalities, such as the alignment of gene expression with (surface) protein expression. In this context, protein features are sparse in the sense that only a limited number of proteins are measured, and the features across modalities exhibit weak linkages, thus constructing an effective guidance graph is particularly challenging.

In this work, we propose BiCLUM, a novel multi-omics integration method that simultaneously enforces cell-level and feature-level alignment across modalities to integrate gene expression (scRNA-seq) with chromatin accessibility (scATAC-seq) or protein expression (CITE-seq). BiCLUM first transforms multi-omics data into a common feature space using prior biological knowledge, such as genepeak or gene-protein relationships, for integrating gene expression with chromatin accessibility or gene expression with protein expression, respectively. It then learns cell and feature embeddings simultaneously through a bilateral contrastive learning framework, incorporating both celllevel and feature-level contrastive losses. For cell-level alignment, BiCLUM utilizes the Mutual Nearest Neighbors (MNN) method to establish correspondences between cells based on their gene expression levels across different modalities. For feature-level alignment, BiCLUM leverages a one-to-one correspondence between shared features, ensuring consistency across datasets. By utilizing these correspondences, BiCLUM employs bilateral contrastive learning to derive a low-dimensional representation of cells, ensuring that similar cell types are clustered together while maintaining distinct separations for different cell types. BiCLUM is applied to integrate gene expression with chromatin accessibility using four scRNA-ATAC-seq multi-omic datasets and a CITE-seq dataset. While pairing information is available for some datasets, it is only used for quantitative evaluation. Extensive benchmarking results show that BiCLUM outperforms other state-of-theart approaches, achieving superior quantitative metrics and visualizations. Furthermore, in downstream biological analysis, BiCLUM demonstrates superior or comparable integration performance compared to existing methods, highlighting its ability to preserve biological characteristics and reveal meaningful biological insights. This robustness in practical applications underscores the potential of BiCLUM for multi-omics data integration.

## Methods

### Problem statement

Given two single-cell datasets, 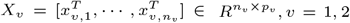, where *n*_*v*_ and *p*_*v*_ represent the number of cells and the dimension of features in the *v*-th modality. Direct integration of *X*_1_ and *X*_2_ is challenging not only due to different cells being assayed but also mismatched feature spaces. To address this, we propose a new method that transforms the datasets from the two modalities into a unified feature space.

We address two integration scenarios within a unified framework: the integration of gene expression with chromatin accessibility and the integration of gene expression with protein expression. We denote the singlecell gene expression matrix as *X*_1_ and the chromatin accessibility or protein expression matrix as *X*_2_.

### BiCLUM model

BiCLUM first transforms *X*_2_ to 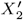 using prior biological knowledge, ensuring that both *X*_1_ and 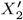 lie within a unified feature space. This transformation leverages prior biological knowledge, such as gene-peak or geneprotein associations, to establish corresponding cell and feature pairs across modalities. Subsequently, the method learns their embeddings simultaneously through a bilateral contrastive learning framework that incorporates both celllevel and feature-level alignment. A contrastive learning method minimizes the distances between representations of similar points (positive pairs) and maximizes the distances between representations of different points (negative pairs). In our method, we define two types of positive pairs in a bilateral way: mutual nearest neighbor (MNN) pairs between cells, and feature pairs between the two modalities. By leveraging these positive pairs in our bilateral contrastive learning model, we could simultaneously align similar cell types and features from both modalities, improving the quality of integration.

Specifically, the BiCLUM method mainly involves three steps: (1) data transformation, (2) cell pairs and feature pairs construction, and (3) bilateral contrastive learning. These steps are illustrated in Fig. 1.

**Fig. 1.**
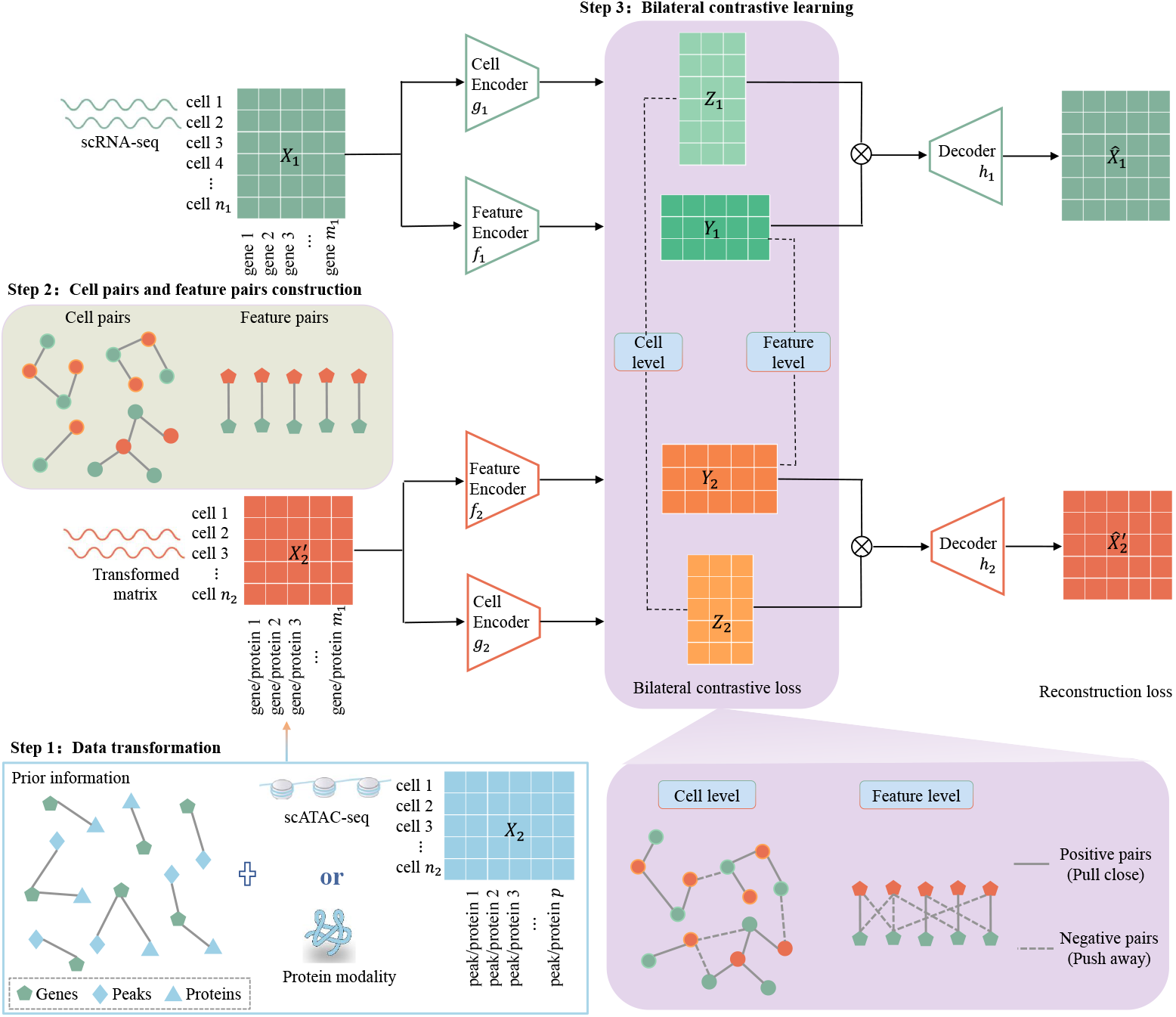
Overview of BiCLUM for unpaired integration of scRNA-seq (*X*_1_) and other modalities such as scATAC-seq or protein data (*X*_2_). The method involves three key steps: (1) step 1: data transformation. Firstly transforming the data from other modalities to obtain a transformed matrix, 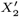.For scATAC-seq data, transforming the modality data into an inferred gene activity score matrix, ensuring that both *X*_1_ and 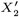 share the same set of features. For the protein modality, the 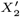 includes only proteins encoded by some of the genes present in *X*_1_. (2) step 2: cell pairs and feature pairs construction. The cell level pairs and the feature level pairs across modalities can be constructed based on *X*_1_ and 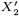.(3) step 3: bilateral contrastive learning. Separate encoders are employed for learning the cell-level (*g*_*v*_, *v* = 1, 2) representations and feature-level (*f*_*v*_, *v* = 1, 2) representations for each modality. The bilateral contrastive learning is to enforce alignment at both the cell and feature levels across modalities. Finally, the latent cell and feature embeddings are passed through modality-specific decoders (*h*_*v*_, *v* = 1, 2) to reconstruct the original features from both modalities.

#### Step 1: Data transformation

Transforming non-RNA modalities to align with the scRNA-seq feature space forms the basis to construct the positive pairs; for either cell pairs or feature pairs, the transformation requires prior information.

To integrate scRNA-seq and scATAC-seq datasets, we first transform scATAC-seq data into a gene activity score matrix using existing methods. These scores estimate gene expression potential by linking chromatin accessibility to genes through regulatory elements. This results in a matrix sharing the same feature space (genes) as scRNAseq data, enabling direct gene pairing for integration. Several methods, including ArchR [23], Cicero [42], cisTopic [7], SnapATAC [30], scATAC-pro [55], Signac [54], and SnapATAC2 [44], have been developed to transform scATAC-seq datasets into gene activity score matrices for downstream analysis, such as inferring cell types. These methods leverage chromatin accessibility data to infer gene expression potential, focusing on the accessibility of gene regulatory regions like promoters and enhancers. The primary difference among them lies in the calculation and weighting of gene region accessibility. In our work, we primarily use ArchR [23] and Signac [54] for data transformation. Detailed descriptions of these methods are provided in the supplementary section 1.2.

The transformation from single-cell protein data to scRNA data is more straightforward, as it involves directly matching proteins to their corresponding encoding genes, leveraging their one-to-one correspondence. By utilizing known biological associations, such as gene-protein relationships or interactions within cellular pathways, we ensure that both modalities are integrated in a biologically relevant manner.

After transformation, we then use a structured preprocessing pipeline that includes several key steps: identifying highly variable genes (HVGs), cell normalization, log transformation, z-score normalization with truncated values, and principal component analysis (PCA). The detailed preprocessing procedure is shown in section 1.3 of the supplementary materials.

#### Step 2: Cell pairs and feature pairs construction

Mutual Nearest Neighbors (MNN) have been proposed [25] and widely used in single-cell RNA-seq integration methods, such as Seurat3 [52] and scDML [60], demonstrating its effectiveness in aligning datasets across modalities. In this study, we construct MNN pairs to integrate different modalities. Cell *i* and cell *j* are considered to form an MNN pair if they are mutually nearest neighbors in their respective modalities. That is, the set of MNN pairs between the two modalities is defined as

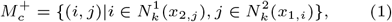

where *x*_*v*,*i*_ represents the *i*-th cell in the *v*-th modality, and 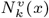 represents the *k*-nearest neighbors of *x* in the *v*-th modality under cosine distances between the cells. For scRNA and gene activity score matrices, we construct MNN pairs based on cosine distances calculated from cross-modal data featuring highly variable genes (HVGs). For scRNA and single-cell protein data, MNN pairs are constructed using scRNA and protein data, retaining only the features present in the gene-protein pairs.

For the construction of feature pairs, as described in step 1, we combine the HVGs from both scRNA and gene activity score matrices to form gene pairs. Our method focuses exclusively on one-to-one correspondences between these gene pairs, excluding co-expressed gene pairs. When integrating scRNA with single-cell protein data, the feature pairs are constructed by matching genes with their corresponding encoded proteins. We define these constructed feature pairs as 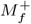.

#### Step 3: Bilateral contrastive learning

Our bilateral contrastive learning is integrated into autoencoder frameworks, ensuring simultaneous alignment of cells and features within low-dimensional embedding spaces. The data *X*_1_ and 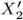 should be well reconstructed through the autoencoders, and meanwhile, the constructed cell pairs and feature pairs should be similar in the embedding spaces. For simplicity, we will use the notation *X*_2_ in place of 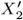 without causing any ambiguity.

Autoencoders are applied to each modality to generate cell embeddings and feature embeddings. Specifically, for each modality *v*, we use distinct cell encoders *g*_*v*_ and feature encoders *f*_*v*_ to learn the cell embeddings 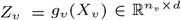 and feature embeddings 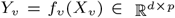 where *d* is the latent dimension and *p* is the number of features. The encoder networks, *f*_*v*_ and *g*_*v*_, are parameterized by neural networks with two hidden layers. *X*_*v*_ is then reconstructed by product of the cell and feature embeddings. A softplus function *h*_*v*_ is applied to this product for nonlinear transformation, resulting in the final reconstructed matrix 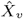:

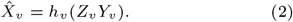

The reconstructed high-dimensional matrix 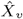 should closely approximate the original input matrix *X*_*v*_. To ensure this, we minimize the reconstruction error by constraining the latent cell and feature embeddings of each modality, using the loss function:

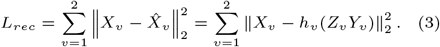

This process ensures that the embeddings capture the essential structure and features of the original data, facilitating effective integration across modalities.

Bilateral contrastive learning is introduced to ensure that the embeddings of the constructed cell pairs and feature pairs are close in the latent space. Cell-level contrastive learning ensures that mutual nearest neighbor (MNN) cell pairs are more closely aligned than other cell pairs, while feature-level contrastive learning ensures that feature pairs are more closely aligned compared to unrelated feature pairs.

For cell-level alignment, we treat the constructed MNN cell pairs in 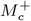 as positive pairs for contrastive learning, while all other cell pairs were regarded as negative pairs, with the set of negative pairs defined as 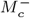.The pairwise contrastive InfoNCE loss is then defined by the following equation:

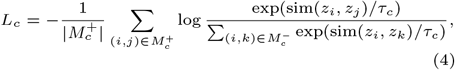

where sim(*z*_*i*_, *z*_*j*_) represents the cosine similarity between the embeddings *z*_*i*_ and *z*_*j*_, *τ*_*c*_ is a temperature constant, and | *·* | represents the number of elements in the set.

Similarly, for feature-level alignment, we treat the constructed feature pairs in 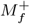 as positive pairs and all other pairs as negative pairs, with the set of negative pairs defined as 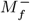.The contrastive loss for feature-level alignment is then defined as:

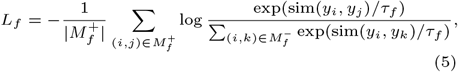

where *τ*_*f*_ is a temperature constant.

Overall, the total loss integrates three components: the reconstruction loss, the contrastive loss for MNN cell pairs across modalities, and the contrastive loss for feature pairs between the modalities,

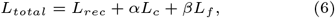

where *α* and *β* are the trade-off parameters to balance the contributions of the different components of the total loss. To streamline training, the process is divided into multiple batches. Based on our experimental results, the parameters *αL*_*c*_ and *βL*_*f*_ are typically set to larger values, with default values of 1*e*4, while the temperature constants*τ*_*c*_ and *τ*_*f*_ are set to 0.5. By employing these steps, we ensure a robust and biologically informed integration of the scRNA-seq and gene activity score matrices or protein data, enhancing the alignment and interpretability of the integrated data.

### Visualization

To assess multi-omics integration methods, we employed several visualization techniques to evaluate performance. First, we used UMAP [36] to project the integrated data into a 2D space, coloring cells by modality or cell type. Although UMAP can intuitively reveal the uniformity of omics data distribution and the separation between cell types, it is important to note that UMAP should not be interpreted as a definitive measure of biological separations. We regard these visualizations as preliminary insights, with further analyses, such as clustering and quantitative assessments, providing more reliable evidence of biological distinctions.

Additionally, we generated confusion matrices to compare predicted clusters with ground truth cell type labels across different methods. Specifically, we applied the Leiden algorithm to cluster cells based on the latent embedding for each method. The number of clusters was set to match the known annotations. Clustering performance was evaluated using accuracy (ACC), comparing predicted clusters to ground truth cell type annotations. To align predicted clusters with cell type labels, we employed the Hungarian algorithm, which minimizes assignment costs to ensure optimal correspondence. In the confusion matrices, diagonal elements represent correctly classified cells, while offdiagonal elements indicate misclassifications. A higher concentration of values along the diagonal suggests better clustering resolution, reflecting the method’s ability to accurately separate cell types.

To further evaluate biological consistency, we conducted trajectory analysis using PAGA [57] graphs. These graphs illustrate differentiation relationships and transition paths between cell types, with edge thickness representing the strength of inferred connections. Thicker edges suggest stronger potential transitions, and greater alignment with known differentiation trajectories indicates better preservation of biological structures.

### Evaluation metrics

Inspired by the previous related work [11, 49] and a benchmarking study [58], we selected four quantitative metrics for integration evaluation, which are omics mixing, cell type conservation, label transfer accuracy and fraction of samples closer than the true match (FOSCTTM).

#### Omics Mixing

- **Graph Connectivity (GC)** evaluates the degree of omics mixing by measuring the largest connected component (LCC) within a k-nearest neighbor (kNN) graph for each cell type. A higher GC score indicates stronger connectivity across omics layers, reflecting better integration.

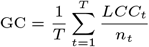

where *LCC*_*t*_ represents the number of cells in the largest connected component for cell type *t, n*_*t*_ is the total number of cells of type *t*, and *T* denotes the total number of cell types. The GC score ranges from 0 to 1, with higher values indicating improved omics integration.
- **Seurat Alignment Score (SAS)** evaluates omics integration by measuring the proportion of k-nearest neighbors from the same omics layer for each cell.

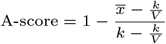

where *k* is the number of neighbors, *V* is the number of omics, and 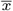 is the average fraction of same-omics neighbors across all cells. SAS ranges from 0 to 1, with higher values indicating better alignment.
- **Average Silhouette Width across Omics (ASWO)** quantifies omics mixing by computing the silhouette width for each cell based on omics layers. The score is defined as:

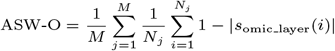

where *s*_omic layer_(*i*) is the silhouette width of cell *i* with respect to its omics layer. Specifically, it measures how well cell *i* is grouped with other cells from the same cell type and omics layer compared to cells from other omics layers within the same cell type. *N*_*j*_ is the number of cells in cell type *j*, and *M* is the total number of cell types. ASW-O ranges from 0 to 1, with higher values indicating better mixing of omics layers across cells.

Omics mixing quantifies the degree of integration across omics layers by aggregating multiple evaluation metrics. A higher omics mixing score indicates better integration and improved consistency between modalities. It is computed as:

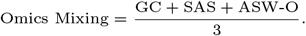

#### Cell Type Conservation

- **Mean Average Precision (MAP)** evaluates cell type resolution by measuring how well a cell’s k-nearest neighbors (kNN) match its true cell type. For each cell, the average precision (AP) is computed as the mean precision at each correctly matched neighbor. MAP is then obtained by averaging AP across all cells:

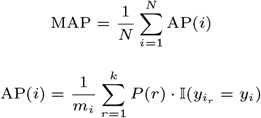

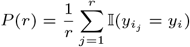

where *k* is the number of neighbors, *y*_*i*_ is the true cell type of cell 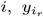 is the cell type of its *r*-th nearest neighbor, and 𝟙(*·*) is an indicator function. *m*_*i*_ is the total number of correctly matched neighbors for cell *i*. MAP ranges from 0 to 1, with higher values indicating better cell type resolution.
- **Average Silhouette Width (ASW)** assesses cell type resolution by measuring how well each cell is clustered with others of the same type. The overall ASW is then obtained by averaging *s*(*i*) across all cells:

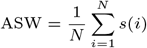

where *s*(*i*) is the silhouette width for a cell *i*, ASW ranges from -1 to 1, with higher values indicating better separation between cell types and stronger clustering.
- **Normalized Mutual Information (NMI)** evaluates cell type resolution by quantifying the agreement between predicted and true cell type labels. It is defined as:

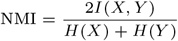

where *X* and *Y* represent the predicted and true cell type label distributions, *I*(*X, Y*) is the mutual information between them, and *H*(*X*) and *H*(*Y*) are their respective entropies. NMI ranges from 0 to 1, with higher values indicating better alignment between predicted and true labels.

The cell type conservation metric evaluates how well the integrated data preserve cell type identities. A higher score indicates better retention of biological distinctions.

It is computed as:

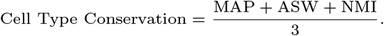

#### Label Transfer Accuracy

- **Label Transfer Accuracy (LTA)** assesses integration quality in transfer learning by evaluating how well cell type labels transfer between omics. One omics dataset is used as the training set, while others serve as test sets. A classifier trained on the training set predicts cell types in the test set, and LTA is defined as the classification accuracy:

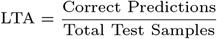 Higher LTA values indicate better cross-omics alignment. By default, a k-NN classifier with *k* = 5 is used, the largest dataset selected as the training set.

#### FOSCTTM

- **Fraction of Samples Closer Than the True Match (FOSCTTM)** quantifies how well predicted cell correspondences align with the ground truth. For each cell in one omics, distances to all cells in the other omics are computed, and the fraction of cells closer than its true match is calculated as:

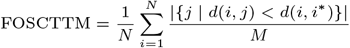

where *d*(*i, j*) is the distance between cell *i* in one omics and cell *j* in the other, *i*^*^ is the true matched cell of *i*, and *M* is the total number of cells in the second omics. FOSCTTM ranges from 0 to 1, with lower values indicating better alignment.

## Results

We evaluated our method on five real multi-omics datasets: BMMC and PBMC (paired), which are paired multi-omics data with scRNA and scATAC modalities; Kidney, an unpaired dataset with snRNA and snATAC modalities; PBMC (unpaired), which includes scRNA and scATAC data; and BMCITE, a paired CITE-seq dataset that simultaneously measures scRNA and protein modalities. While some of the datasets are paired, we treated the paired data as unpaired, using the pairing information solely for quantitative evaluation purposes. A summary of these datasets, including the information of raw data and the dimensionality of the transformed matrices, is provided in Table 1.

**Table 1.** Summary of datasets used to evaluate the method. Each dataset includes pairing information and cell type annotations as ground truth for benchmarking.

| Dataset | Modality | Sample Details | Source | #Cells | #Features | Ground Truth |
| --- | --- | --- | --- | --- | --- | --- |
| **BMMC** | scRNA + scATAC | Bone marrow mononuclear cells | GSE194122 [29, 33] | 6,224 | 13,431 genes, 116,490 peaks, 19,607 transformed features, 11,141 gene-transformed feature pairs | Paired data; cell type annotation |
| **Kidney** | snRNA + snATAC | Human adult kidney cortex samples | GSE151302 [38] | 19,985 (snRNA), 27,034 (snATAC) | 33,538 genes, 196,113 peaks, 24,919 transformed features, 21,277 gene-transformed feature pairs | Cell type annotation |
| **PBMC (paired)** | scRNA + scATAC | Peripheral blood mononuclear cells | 10X Genomics[1] | 9,631 | 29,095 genes, 107,194 peaks, 24,919 transformed features, 19,016 gene-transformed feature pairs | Paired data; cell type annotation |
| **PBMC (unpaired)** | scRNA + scATAC | Peripheral blood mononuclear cells | [55, 12] | 1,985(scRNA), 1,919(scATAC) | 2,000 genes, 40,850 peaks, 1,931 transformed features, 1,735 gene-transformed feature pairs | Cell type annotation |
| **BMCITE** | scRNA + protein | Bone marrow mononuclear cells | GSE194122 [29, 33] | 10,205 (s1d1/s1d2) or 16,451 (s1d2/s3d7) | 13,953 genes, 134 proteins, 134 transformed features, 95 gene-transformed feature pairs | Paired data; cell type annotation |

We compared our method with 13 state-of-the-art methods, categorized into three groups: six methods that integrate raw multi-omics data through nonlinear manifold alignment, five methods that use prior genomic information to transform features from other modalities into a gene activity score matrix, which shares a common feature space with scRNA-seq data, and two methods that leverage regulatory network information to construct a bridge between features across modalities. These methods are listed in Table 2, with detailed descriptions provided in Section 1.1 of the supplementary materials.

**Table 2.** The compared single cell multi-omics integration methods.

| Input | Methods | Software |
| --- | --- | --- |
| Raw data | MMD-MA [49] | Python |
|  | UnionCom [11] | Python |
|  | Scot [19] | Python |
|  | Pamona [13] | Python |
|  | JointMDS [16] | Python |
|  | scTopoGAN [48] | Python |
| Raw scRNA-seq+gene activity score matrix | Seurat3 [52] | R |
|  | LIGER [32] | R |
|  | uniPort [12] | Python |
|  | MultiMAP [27] | Python |
|  | Bi-CCA [20] | R |
| Raw data + regulatory network | scDART [61] | Python |
|  | GLUE [14] | Python |

To ensure a fair comparison, we applied the same gene activity score transformation across all methods that required this preprocessing step. Specifically, we used either ArchR or Signac to generate gene activity score matrices from scATAC-seq data. The selected transformation method and parameter settings for each dataset of our BiCLUM method are summarized in Table S1.

### BiCLUM achieves superior quantitative metrics for the integration of scRNA and scATAC data in BMMC data

We first evaluated BiCLUM on the integration of scRNA and scATAC data using the BMMC dataset, comparing its performance with raw data and 13 state-of-theart methods. The evaluation included visualization of integration results and quantitative assessments.

In Fig. 2A, we present UMAP visualizations of raw, unintegrated data and embeddings produced by BiCLUM. Embeddings from the various methods are visualized in Fig. S1. In all plots, cells are color-coded based on their omics type and cell type. The results demonstrate that GLUE, Seurat, and BiCLUM effectively integrates the two modalities, with most cells of the same type clustering together and distinct cell types being separated. In contrast, the other compared methods either fail to mix the omics data effectively or do not achieve good separation of different cell types.

**Fig. 2.**
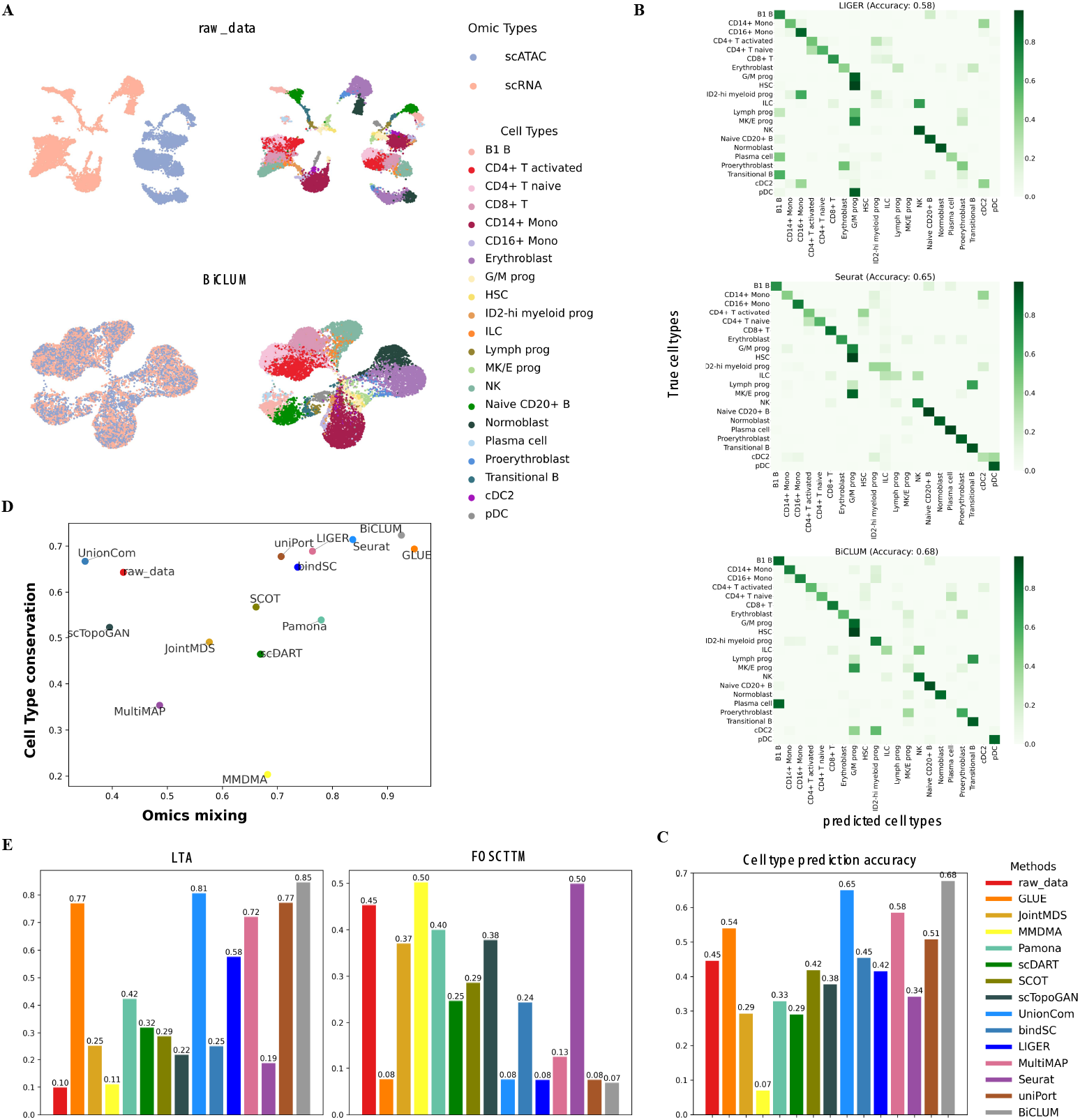
Integrated results of BMMC data. (A) UMAP visualization of the embeddings for the raw data and BiCLUM method with cells colored based on omics types and cell types, respectively. (B) Heatmap of the confusion matrix for LIGER, Seurat and BiCLUM, where rows represent the true cell types, columns represent the predicted cell types, and each element *i, j* in the matrix represents the proportion of cells of type *i* that are classified as type *j*. (C) Cell type prediction accuracy values across different integration methods. (D) Two evaluation metrics of omics mixing and biology conservation for multi-omics integration methods. (E) LTA and FOSCTTM across different integration methods.

To further assess the biological relevance of these embeddings, we performed trajectory analysis using PAGA graphs (Fig. S3). BiCLUM effectively reconstructs key differentiation pathways in hematopoiesis, preserving welldefined transitions from hematopoietic stem cells to progenitors and differentiated cell types. It accurately captures lymphoid and myeloid lineage progressions, erythropoiesis, and dendritic cell development, aligning with known biological hierarchies [31, 2, 59, 15]. In contrast, some comparison methods produce disordered graphs that obscure differentiation trajectories, though GLUE, UnionCom and uniPort successfully capture certain key transitions. These results highlight BiCLUM’s strength in both modality alignment and biological trajectory preservation.

Beyond qualitative visualization and trajectory analysis, we quantitatively evaluated cell type separation using confusion matrices (Fig. 2B and Fig. S2), which compare true and predicted cell types across different methods. Higher values along the diagonal indicate better alignment, with Seurat and BiCLUM demonstrating the most accurate predictions and minimal misclassification. Furthermore, clustering accuracy analysis (Fig. 2C) confirms BiCLUM’s superior performance, achieving the highest overall accuracy of 0.68, surpassing all other methods in distinguishing cell types effectively.

Quantitative performance is summarized in Fig. 2DE. Fig. 2D compares methods based on omic mixing and cell type conservation. BiCLUM achieves superior integration (highest omic mixing) and maintains high cell type conservation, closely following GLUE. Fig. 2E evaluates LTA and FOSCTTM. BiCLUM achieves the highest LTA and the lowest FOSCTTM value, indicating superior alignment of multi-omics data and clustering of similar cell types.

### BiCLUM excels in integrating multi-omics Kidney data and achieves well-mixed omics and distinct separation of cell types

We further evaluated the performance of BiCLUM on the Kidney dataset. As shown in Fig. 3A and Fig. S4, UMAP plots illustrate the integration results for various methods. In the raw data, omics datasets are not well mixed, with most cell types nearly separated, except for certain cell types like CNT and DCT, which are not clearly clustered. BiCLUM demonstrates superior performance by not only achieving effective mixing of the two omics datasets but also ensuring that cells of the same type are well-clustered. In comparison, other methods show varying levels of integration quality. While some methods, such as scDART, GLUE, Seurat, and bindSC, successfully mix scRNA-seq and scATAC-seq data, their cell type separation remains suboptimal. For instance, bindSC exhibits significant mixing of distinct cell types. scDART struggles to separate certain cell types with fewer cells. GLUE fails to distinguish between PODO cells and LEUK cells, as well as between ICA and ICB cells. Seurat encounters difficulties in separating LEUK and ENDO cells, leading to partial overlap between these two types.

**Fig. 3.**
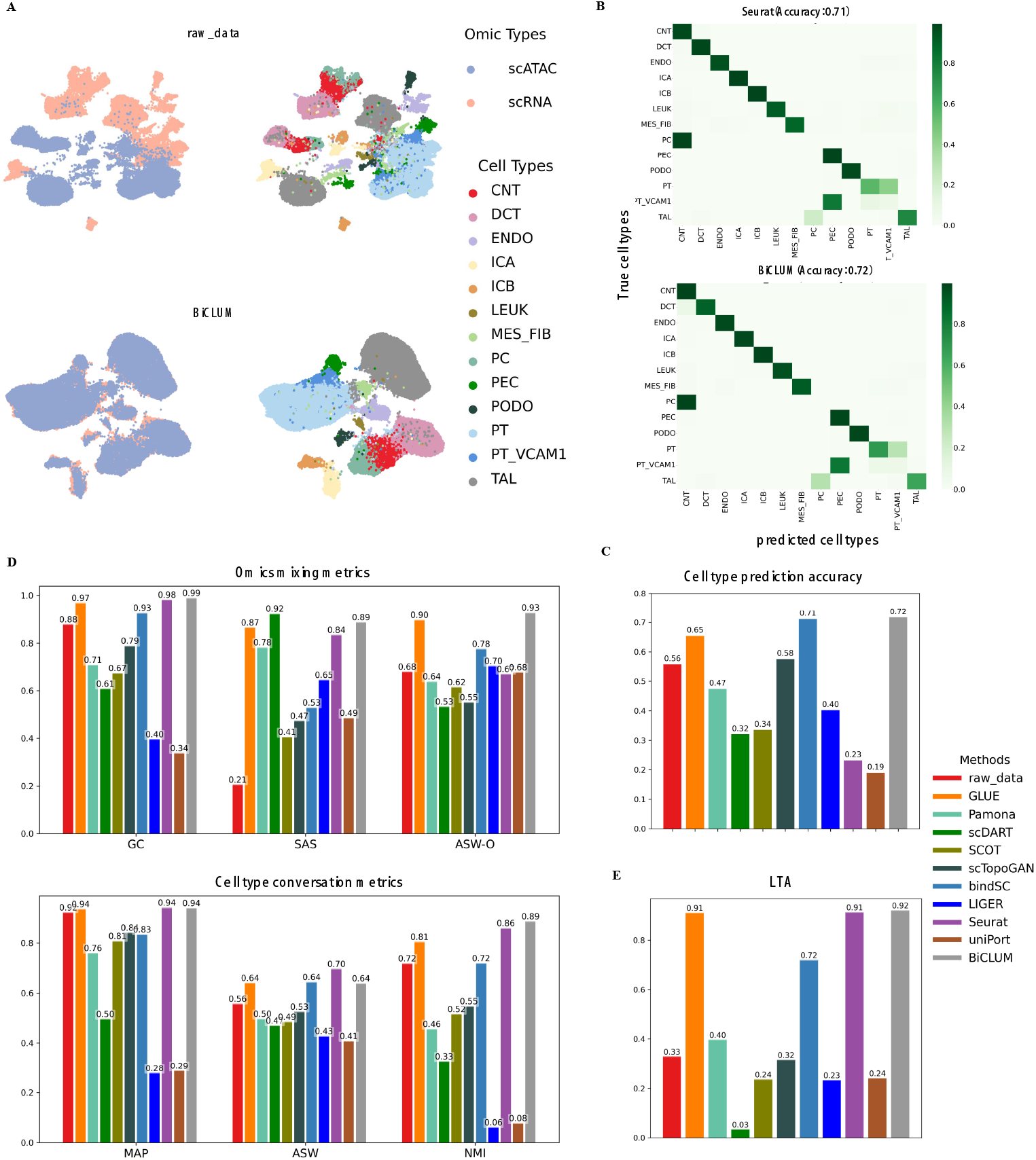
Integrated results of kidney data. (A) UMAP visualization of the embeddings for the raw data and BiCLUM method with cells colored based on omics types and cell types, respectively. (B) Heatmap of the confusion matrix for Seurat and BiCLUM, where rows represent the true cell types, columns represent the predicted cell types, and each element *i, j* in the matrix represents the proportion of cells of type *i* that are classified as type *j*. (C) Cell type prediction accuracy values across different integration methods. (D) All individual metrics belonging to the two evaluation categories, omics mixing (GC, SAS, ASW-O) and biological conservation (MAP, ASW, NMI), in the assessment of multi-omics integration methods. (E) LTA across different integration methods.

Furthermore, the trajectory analysis based on PAGA graphs (Fig. S5) highlights BiCLUM’s ability to preserve biologically meaningful functional relationships among kidney cell types. Unlike methods that produce overly complex or ambiguous trajectories, BiCLUM accurately captures key interactions, such as those among glomerular cells (PT, PEC, PODO) [47, 28], distal nephron components (TAL, DCT, CNT, PC) [56], immune and endothelial cells (LEUK, ENDO, MES-FIB) [6, 17], and intercalated cells (ICA, ICB) [40]. These results demonstrate that BiCLUM not only achieves superior multi-omics integration but also effectively preserves functional relationships, reinforcing its biological interpretability.

Fig. 3B displays the confusion matrices for Seurat and BiCLUM, illustrating their performance in cell type classification. The diagonal elements in the matrix highlight accurate predictions for both methods. Fig. 3C further quantifies clustering accuracy across methods, with BiCLUM achieving the highest accuracy of 0.72. This underscores its effectiveness in maintaining cell type distinctions during integration.

Figure 3D quantifies the performance of different methods across individual metrics associated with omics mixing and cell type conservation. Notably, BiCLUM achieves the highest scores for GC and ASW-O, indicating superior omics integration. In terms of cell type conservation, it attains the highest MAP and NMI values, while the remaining metrics rank second best, further demonstrating its ability to preserve distinct cellular identities. Moreover, Fig. 3E demonstrates that BiCLUM achieves the highest LTA among all methods, further emphasizing its effectiveness in integrating unpaired multiomic data.

### BiCLUM achieves robust multi-omics PBMC integration with well cell type preservation

To further evaluate the performance of BiCLUM, we analyzed two PBMC datasets: a paired PBMC dataset, which contains simultaneous scRNA-seq and scATAC-seq measurements, and an unpaired PBMC dataset, where the scRNA and scATAC modalities are obtained separately. Given the critical role of gene activity score matrices in integrating scRNA and scATAC data, we also assessed the impact of different transformation methods.

#### Paired PBMC Data Analysis

The paired PBMC dataset was also analyzed under the same experimental settings. In Fig. 4A and Fig. S6, the UMAP visualizations illustrate the integration outcomes of various methods. The raw data exhibits poor integration, with scATAC-seq and scRNA-seq measurements forming distinct, non-overlapping batches. Methods such as bindSC, LIGER, Pamona, and scDART demonstrate improved integration, with partial overlap between the two omics datasets, though the results remain suboptimal. In contrast, GLUE, Seurat, and BiCLUM achieve the most refined integration, characterized by well-mixed scATACseq and scRNA-seq data points and distinct clustering for each cell type.

**Fig. 4.**
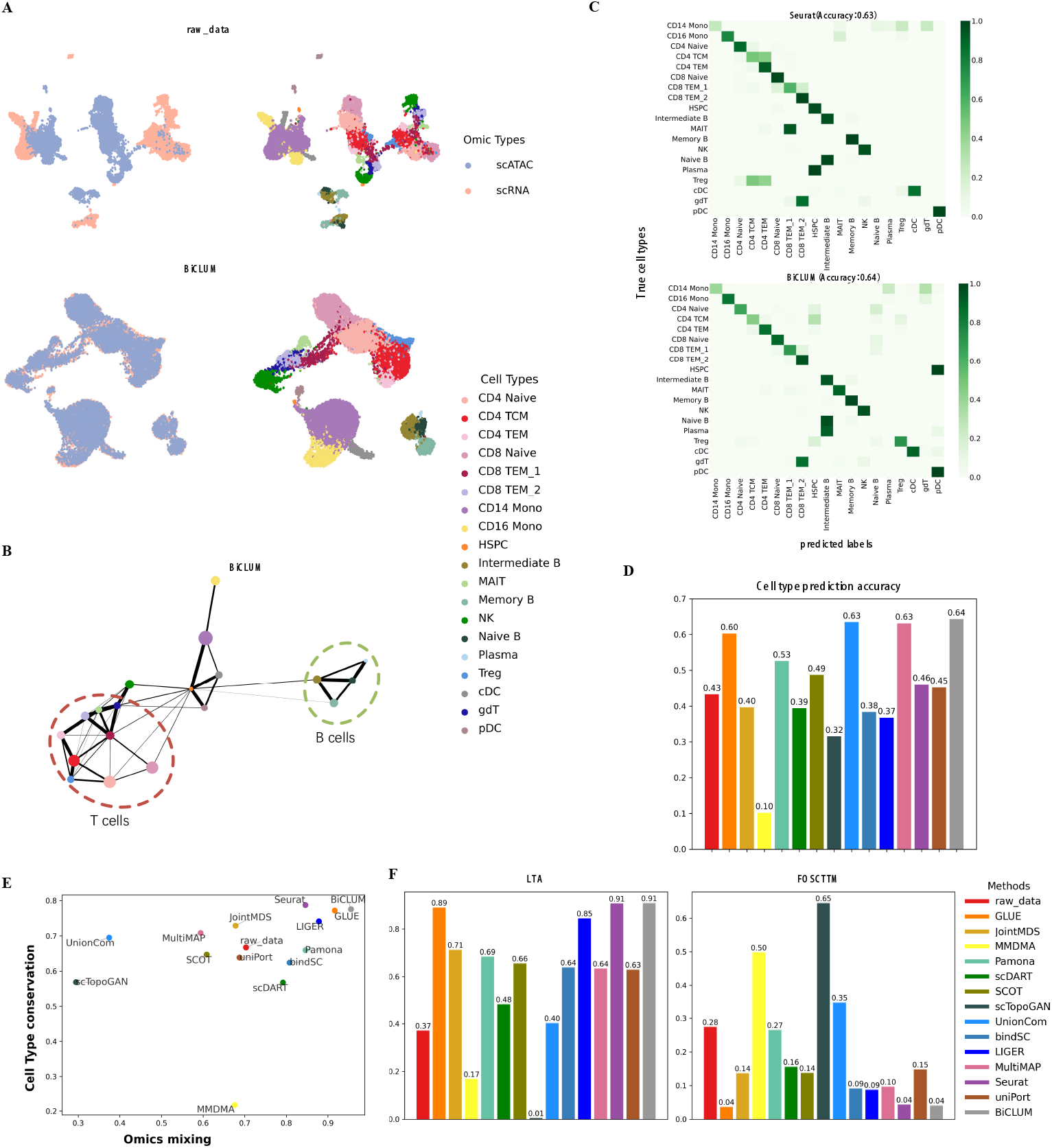
Integrated results of PBMC (paired) data. (A) UMAP visualization of the embeddings for the raw data and BiCLUM method with cells colored based on omics types and cell types, respectively. (B) PAGA trajectory visualizations of the integrated embeddings for BiCLUM method, where each node represents a cell type and the size is proportional to the number of cells in that type. Edges indicate potential lineage relationships, with the thickness representing the degree of connectivity between cell types. Cell types from the same lineage are grouped and highlighted with dotted ovals for clarity. (C) Heatmap of the confusion matrix for Seurat and BiCLUM, where rows represent the true cell types, columns represent the predicted cell types, and each element *i, j* in the matrix represents the proportion of cells of type *i* that are classified as type *j*. (D) Cell type prediction accuracy values across different integration methods. (E) Two evaluation metrics of omics mixing and biology conservation for multi-omics integration methods. (F) LTA and FOSCTTM values of different integration methods.

Fig. 4B displays PAGA graphs of the BiCLUMintegrated results, which effectively highlight the functional and developmental relationships among cell types. BiCLUM captures key differentiation trajectories, such as the differentiation of CD4+ and CD8+ Naïve cells into Treg, TCM, and TEM cells, as well as the differentiation of naive B cells into intermediate B cells, memory B cells, and plasma cells. These observations are in alignment with existing literature [46, 21, 22], thereby validating the biological relevance of BiCLUM’s integration and trajectory inference. Additionally, PAGA graphs for comparison methods are shown in Fig. S7. Notably, with the exception of the MMDMA method, most methods reveal relatively clear trajectory paths. However, methods such as GLUE, MultiMAP, and UnionCOM fail to reflect the role of HSPCs as the origin of all blood cell types, displaying few or weak edges emanating from HSPCs. Furthermore, methods such as bindSC, Seurat, and SCOT misplace plasma cells near HSPCs, leading to biologically implausible connections with multiple cell types, which contradicts their distinct lineage origins. The PAGA graph analysis, coupled with the UMAP clustering shown in Fig. 4A, underscore BiCLUM’s superior performance in integrating multi-omics data and producing biologically meaningful differentiation trajectories.

Fig. 4C compares the confusion matrices for Seurat and BiCLUM. While the accuracy values across the methods are similar, BiCLUM demonstrates a clearer diagonal pattern, indicating more accurate cell type assignments and reduced misclassification. Furthermore, Fig. 4D quantitatively evaluates the accuracy across all methods, with BiCLUM achieving the highest accuracy of 0.64. This underscores its superior ability to preserve cell type identities during integration.

Fig. 4E further quantifies the integration outcomes, while Seurat achieves the highest cell type conservation, its omics mixing value is lower than BiCLUM’s. BiCLUM balances both metrics effectively. Lastly, Fig. 4F shows that BiCLUM achieves high LTA and the lowest FOSCTTM values, indicating superior alignment and integration across omics datasets.

#### Unpaired PBMC Data Analysis

We further evaluated BiCLUM on an unpaired, microfluidic-based PBMC dataset. Unlike the paired PBMC dataset, the transformed gene activity score matrix in this case was generated using the MAESTRO method, as provided by the original study. The UMAP visualizations in Fig. S8A show that GLUE, uniPort, and BiCLUM achieve the best integration, effectively aligning scRNA and scATAC data while maintaining relatively clear cell-type separation. Quantitative metrics in Fig. S8B-C indicate that GLUE achieves the highest performance, followed by BiCLUM, which exhibits slightly lower omics mixing and cell type conservation scores but comparable transfer accuracy.

#### Impact of Gene Activity Score Matrix Transformations

Since BiCLUM relies on the transformed gene activity score matrix for scRNA and scATAC integration, we evaluated the impact of different transformation methods (Fig. S9). We assessed the average Pearson correlation between constructed MNN pairs and the proportion of MNN pairs with matching cell types. The results indicate that ArchR and Signac are the most stable methods, consistently producing high-quality MNN pairs. In contrast, cisTopic and SnapATAC2 generally perform well but occasionally show reduced performance. Cicero, Gene Scoring, and MAESTRO exhibit relatively lower overall performance compared to other methods. Based on these findings, we recommend using ArchR and Signac for optimal gene activity score matrix transformation when applying BiCLUM.

### BiCLUM reveals consistent, biologically relevant patterns and achieves superior quantitative metrics across BMCITE data

Integration of gene expression and protein abundance is a challenging task as the correlation between mRNA and protein levels can be weak, which is due to factors such as post-transcriptional modifications, differences in degradation rates, and other regulatory mechanisms [24]. Despite these challenges, integrating single-cell RNA and protein data offers significant potential to uncover cellular diversity and functional states. For example, integrating transcriptomic and proteomic data has been used to generate comprehensive maps of aging lung tissue, quantifying activity state changes across cell types [4]. Several computational methods, such as Seurat3, bindSC and so on have applied to the integration of scRNA and protein data.

We tested our method on a CITE-seq dataset derived from the same cohort of 12 healthy human donors. For evaluation, we selected three subsets of cells, s1d1, s1d2, and s3d7, from the complete dataset. Specifically, we integrated scRNA and ADT for cells from s1d1 and s1d2, with results reported in Fig. 5, and for cells from s1d2 and s3d7, with results shown in Fig. 6. This design allowed us to evaluate whether our method achieves robust and biologically meaningful integration, ensuring that consistent patterns reflecting true biological insights are identified across datasets without being confounded by data-specific effects. Fig. 5A and Fig. 6A display clustering results for scRNA-seq data (featuring highly variable and protein-coding genes) and protein data (featuring the corresponding gene-encoded proteins). These results demonstrate that while protein data achieve a comparable accuracy value to scRNA data, cells of the same type in the protein dataset are not as tightly clustered. Further analysis revealed 66% and 62% match rate for cells of the same type within the constructed MNN pairs for the two sets respectively. This indicates that the MNN method effectively aligns features across modalities by utilizing local neighborhood information, even in scenarios where global correlations are weak. This observation is consistent with prior studies showing that RNA and protein measurements capture complementary aspects of cellular biology, representing distinct yet interrelated regulatory layers [51, 53].

**Fig. 5.**
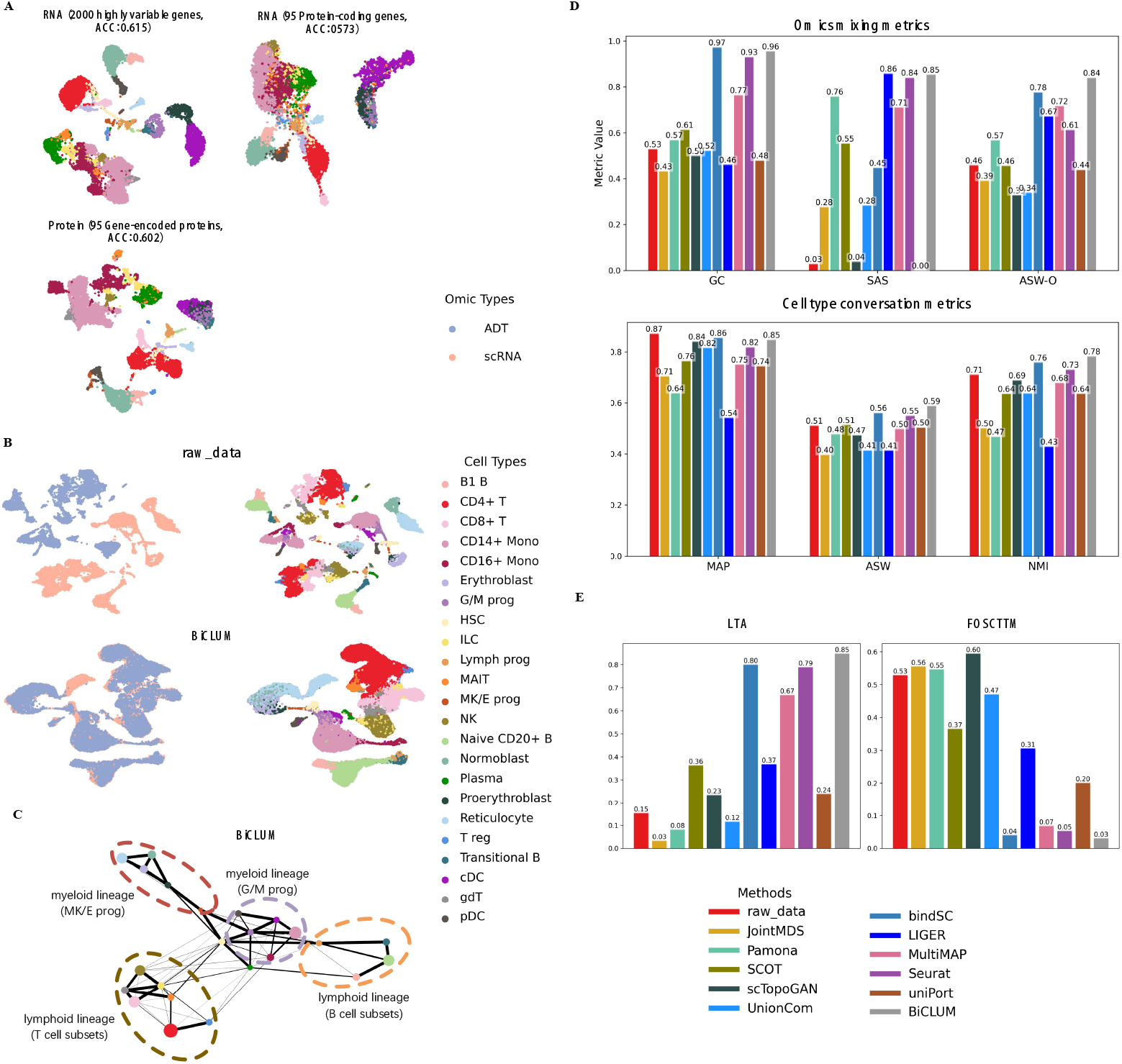
Integrated results of BMCITE data for the two sets, s1d1 and s1d2. (A) UMAP visualization of scRNA-seq data (featuring highly variable and protein-coding genes) and protein data (featuring the corresponding gene-encoded proteins) along with corresponding clustering accuracies. (B) UMAP visualization of the embeddings for the raw data and BiCLUM method, with cells colored based on omics types and cell types, respectively. (C) PAGA trajectory visualizations of the integrated embeddings for the BiCLUM method, where each node represents a cell type, with the size proportional to the number of cells in that type. Edges indicate potential lineage relationships, with thickness representing the degree of connectivity between cell types. Cell types from the same lineage are grouped and highlighted with dotted ovals for clarity. (D) All individual metrics belonging to the two evaluation categories, omics mixing (GC, SAS, ASW-O) and biological conservation (MAP, ASW, NMI), in the assessment of multi-omics integration methods. (E) LTA and FOSCTTM values for different integration methods.

**Fig. 6.**
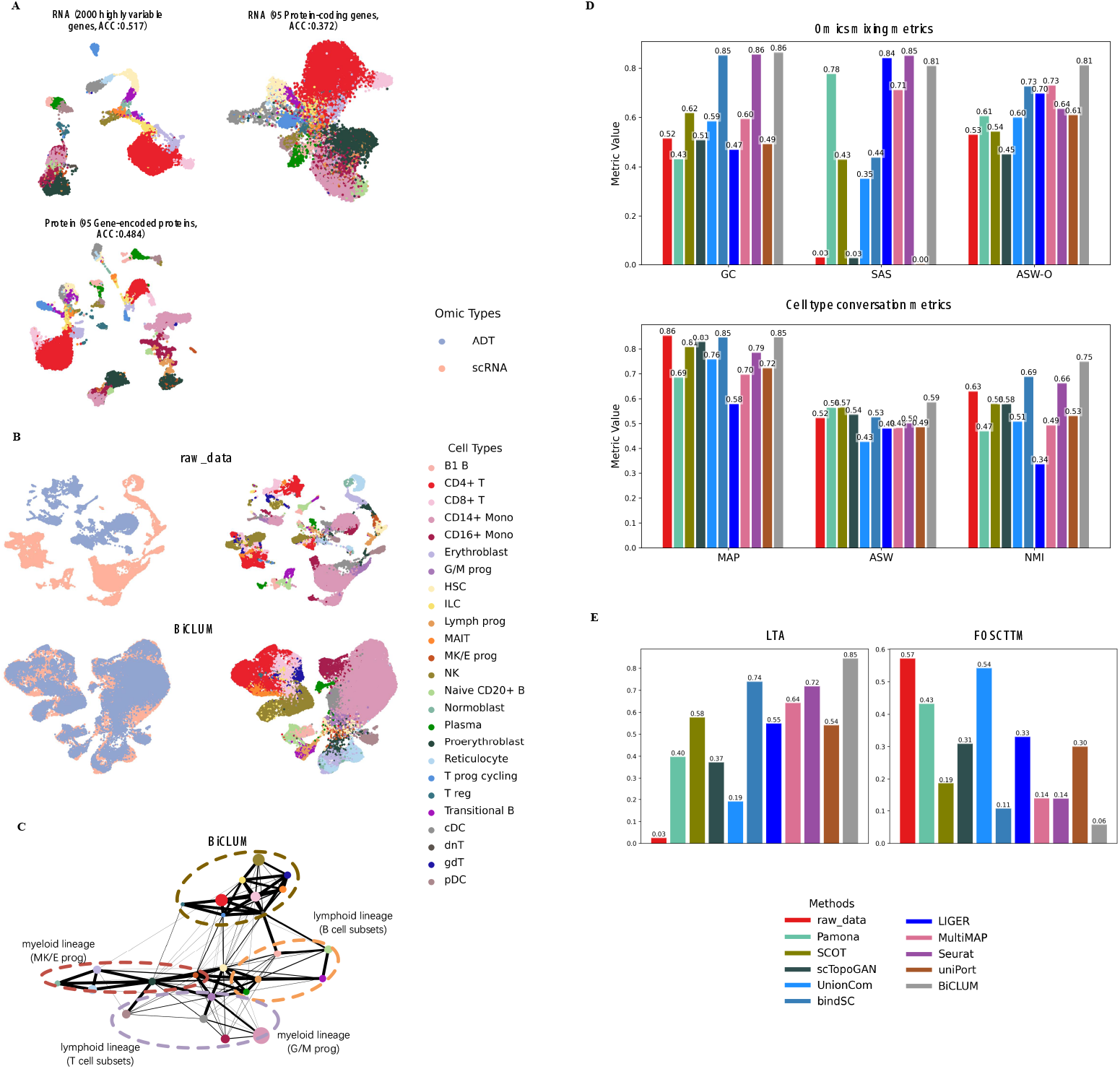
Integrated results of BMCITE data for the two sets, s1d2 and s3d7. (A) UMAP visualization of scRNA-seq data (featuring highly variable and protein-coding genes) and protein data (featuring the corresponding gene-encoded proteins) along with corresponding clustering accuracies. (B) UMAP visualization of the embeddings for the raw data and BiCLUM method, with cells colored based on omics types and cell types, respectively. (C) PAGA trajectory visualizations of the integrated embeddings for the BiCLUM method, where each node represents a cell type, with the size proportional to the number of cells in that type. Edges indicate potential lineage relationships, with thickness representing the degree of connectivity between cell types. Cell types from the same lineage are grouped and highlighted with dotted ovals for clarity. (D) All individual metrics belonging to the two evaluation categories, omics mixing (GC, SAS, ASW-O) and biological conservation (MAP, ASW, NMI), in the assessment of multi-omics integration methods. (E) LTA and FOSCTTM values for different integration methods.

We then reported the UMAP visualizations of integration results for different methods on the two constructed datasets. In Fig. 5B and Fig. 6B, we presented the UMAP visualizations of the raw data alongside the embeddings produced by BiCLUM. The UMAP visualizations of the corresponding embeddings after integration by the comparison methods are shown in Fig. S10 and Fig. S11. From these visualizations, it can be seen that BiCLUM effectively integrates cells from the two modalities, achieving well-mixed embedding with clear separation between most cell types. In contrast, the compared methods demonstrate varying levels of integration but exhibit notable limitations. For instance, while bindSC and Seurat achieve reasonable integration of the two modalities, their performance falls short in certain aspects, such as the inadequate alignment of reticulocyte cell types across modalities. Other methods either fail to fully integrate the two data modalities or struggle to maintain clear separation between cell types, resulting in overlapping and poorly defined clusters.

Furthermore, Fig. 5C and Fig. 6C illustrate the PAGA graphs for both datasets, revealing consistent and biologically relevant differentiation patterns. BiCLUM accurately reconstructs hematopoietic trajectories, capturing well-defined transitions from granulocyte-monocyte progenitors (G/M Progs), megakaryocyte-erythroid progenitors (MK/E Progs), and lymphoid progenitors (Lymph Progs). The central positioning of HSCs and the structured progression of B cells, T cells, monocytes, and erythroid cells align with established hematopoietic hierarchies [39, 9, 3, 50], demonstrating the robustness of BiCLUM’s integration.

For both datasets, we further assessed the integration performance of different methods using multiple quantitative metrics evaluating omics mixing and cell type preservation (Fig. 5E and Fig. 6E). BiCLUM consistently ranked first or second across nearly all metrics, demonstrating its ability to achieve a well-balanced integration that preserves both omics consistency and biological identity. Furthermore, as shown in Fig. 5F and Fig. 6F, BiCLUM achieved the highest LTA among all methods while maintaining competitive FOSCTTM scores. These results underscore BiCLUM’s effectiveness in integrating multimodal data, ensuring both accurate data alignment and biologically meaningful structure retention.

By evaluating the integration results across multiple subsets of cells, we have demonstrated that the patterns observed are not only robust but also reproducible, underscoring the method’s capacity to generalize across different datasets. This consistency further affirms BiCLUM’s ability to capture true biological relationships.

## Discussion and Conclusion

In this work, we introduce BiCLUM, a novel method for integrating unpaired multi-omics data by bilaterally aligning both cells and features across modalities.

Our approach combines the concept of mutual nearest neighbors (MNN) for cell-level alignment and known genomic information for feature-level alignment across modalities. Using these correspondences, we form positive and negative pairs for contrastive learning, which allows for alignment at both the cell and feature levels. By adopting this strategy, BiCLUM achieves that different modalities are well-mixed while preserving the separation between different cell types.

Through comprehensive evaluation on four scRNAseq and scATAC-seq datasets and one CITE-seq dataset, we demonstrate that BiCLUM compares favourably to existing integration methods, achieving strong performance in both visualization and quantitative metrics. Notably, BiCLUM excels in preserving biologically meaningful relationships across modalities, such as how PAGA graphs are consistent with existing literature. Its cell type preservation consistently outperforms other methods based on quantitative metrics, making it a useful tool for downstream biological analysis. Furthermore, our results indicate that using ArchR and Signac produces higher-quality gene activity score matrices and more accurate MNN pairs, leading to improved integration performance. We therefore recommend these methods for gene activity score matrix transformation when applying BiCLUM to scRNA and scATAC data.

Despite these promising results, there are several areas for further improvement. While BiCLUM assumes a straightforward one-to-one correspondence between features across modalities, this may not be sufficient for more complex datasets, where co-expressed or indirectly related genes play a significant role. Additionally, our current approach focuses on integrating two modalities, but the integration of three or more modalities, or modalities with limited or no prior biological information, presents additional challenges. Future work could explore more flexible alignment strategies that better account for complex relationships between modalities. Furthermore, incorporating additional biological constraints or adopting more advanced representation methods, such as graphbased techniques, could enhance the integration quality and provide deeper insights into the multimodal data. Overall, leveraging established biological knowledge in combination with advanced machine learning techniques represents a promising direction for the integration of multimodal single-cell data. Approaches that combine biological constraints, domain-specific insights, and machine learning have great potential to enhance robustness and expand applicability across a wider range of biological contexts.

### Key Points

- Unpaired Single-Cell Multi-Omics Data Integration: BiCLUM effectively integrates diverse unpaired multi-omics datasets by aligning both cell-level and feature-level information across modalities, addressing the challenge of integrating different types of omics data.
- Bilateral Contrastive Learning Framework: BiCLUM introduces a novel bilateral contrastive learning approach, leveraging mutual nearest neighbors (MNNs) for cell alignment and integrating prior genomic knowledge for feature alignment. This approach enables the construction of positive pairs for contrastive learning and facilitates data integration.
- Performance: BiCLUM compares favourably to existing integration methods across multiple datasets, excelling in both visualization and quantitative metrics. This highlights its effectiveness in integrating unpaired multi-omics data with high-quality results.
- Application Potential for Biological Analysis: BiCLUM offers robust downstream analysis capabilities, including cell type identification and trajectory exploration, making it a potentially powerful tool for generating meaningful biological insights and advancing our understanding of cellular processes.

## Supplementary Data

Supplementary data are available online at *Briefings in Bioinformatics*.

## Data availability

The source code for BiCLUM and for reproducing the results are publicly available on GitHub (ht tps://github.com/LiminLi-xjtu/BiCLUM and https://github.com/LiminLi-xjtu/BiCLUM_test), and the datasets used for evaluations can be accessed on Zenodo (https://zenodo.org/records/14506611). For the original multi-omics datasets, Kidney dataset was downloaded from https://www.ncbi.nlm.nih.gov/geo/query/acc.cgi?acc=GSE151302. PBMC (paired) dataset was downloaded from https://www.10xgenomics.com/resources/datasets/pbmc-from-a-healthy-donor-granulocytes-removed-through-cell-sorting-3-k-1-standard-2-0-0. PBMC (unpaired) dataset was downloaded from https://github.com/liulab-dfci/MAESTRO/tree/master/data. BMCITE and BMMC datasets were downloaded from https://www.ncbi.nlm.nih.gov/geo/query/acc.cgi?acc=GSE194122.

## Author Biographies

Yin Guo is currently working toward the PhD degree in Xi’an Jiaotong University, China. Her research interest includes biostatistics.

Izaskun Mallona is a research associate in University of Zurich, Switzerland. Her research interest includes methods development, gene expression regulation, and single-cell technologies.

Mark Robinson is a professor of University of Zurich, Switzerland. His research interest includes biostatistics.

Limin Li is a professor of Xi’an Jiaotong University, China. Her research interests include machine learning and the applications in bioinformatics and biostatistics.

## Acknowledgments

This work was funded by National Natural Science Foundation of China projects under Grant No. 12222115 and 92470106.

