## Supplemental Figure 1-11, Supplemental Table 1 for "BiCLUM: Bilateral Contrastive Learning for Unpaired Single-Cell Multi-Omics Integration"

### 1 Additional materials

#### 1.1 Compared methods

The existing methods can be roughly devided into three types:

- **Learn latent embeddings for original omics layers via nonlinear manifold alignment: MMD-MA [16], UnionCom [2], Scot [7], Pamona [4], Joint-MDS [6], scTopoGAN [15].** UnionCom is a two-step method which first finds the correspondence between the cells from different omics by aligning the corresponding kernels, and then find low-dimensional representations by preserving the correspondence and local structure. MMD-MA integrates cells measured in different ways into a common latent space based on the Maximum Mean Discrepancy (MMD), preserving the structure of the original high-dimensional dataset in the low-dimensional embedding. Joint MDS integrates the multidimensional scaling (MDS) and Wasserstein Procrustes analysis into a joint optimization problem to simultaneously learn the low-dimensional latent embedding of each omic and the correspondence between cells from two different omics. Pamona develops a partial Gromov-Wasserstein optimal transport framework to divide cells in different omics into shared and dataset-specific cells, which can align the cells in a common low-dimensional space while preserving shared and dataset-specific structures. SCOT is also a Gromov-Wasserstein-based optimal transport method to align single-cell multi-omics datasets. The method first constructs a $k$-NN graph for each omic and finds a probabilistic coupling between the cells of each omic to minimize the distance between the graph distance matrices generated by the $k$-NN graph. At last, the coupling matrix is used to perform alignment by projecting one single cell dataset onto another single cell dataset. scTopoGAN uses topological autoencoders to obtain latent representations of each modality separately. A topology-guided Generative Adversarial Network then aligns these latent representations into a common space.

- **Convert multimodality data into one common feature space based on prior knowledge : Seurat3 [17], LIGER [12], uniPort [3], MultiMAP [10],**

**bindSC [8].** Seurat3 is an alignment workflow integrating single-cell multi-omics datasets with CCA amd MNN procedures. LIGER identifies shared and dataset-specific factors through integrative non-negative matrix factorization. uniPort is also a method that also makes use of the optimal transport strategy and the omic-specific features are considered in the method. MultiMAP learns a low-dimensional representation of multimodal data based on the UMAP framework. bindSC also uses CCA method, the integratation includes: the scRNA and GAM datasets, and the scATAC and GAM datasets.

- **Make use of the information of regulatory network during the integration process: scDART [20], GLUE [5].** GLUE learns latent embeddings by constructing a knowledge-guided graph to link features across modalities and employs an adversarial network to align cells. scDART also uses prior knowledge to define a linkage matrix between features across modalities and aligns the datasets through MMD.

#### 1.2 Methods for obtaining gene activity score matrices

Gene activity score can be used to infer the cell types for scATAC-seq datasets. There have been several methods analyse the scATAC-seq datasets by the gene activity score matrices.

- **ArchR [9]** creates a tile matrix based on a user-defined tile size and overlaps these tiles with a user-defined gene window. It calculates the distance from each tile to the gene body or TSS and identifies the tiles within the gene window that don't overlap with other genes. The distance is then converted to a weight using a user-defined accessibility model (default is e(-abs(distance)/5000) + e-1). To account for gene size differences, ArchR applies an inverse gene size weight (1/gene size), scaled linearly from 1 to a user-defined maximum (default: 5). The distance and gene size weights are multiplied by the number of Tn5 insertions in each tile and summed across all tiles within the gene window. This sum gives the gene score, which is normalized to a user-defined constant (default: 10,000).

- **Signac [18]** calculates gene activity scores by determining the number of fragments within genomic regions associated with each gene. The process starts by extracting gene coordinates and expanding them to include an upstream region of 2kb to cover regulatory elements in the promoter region. The FeatureMatrix() function is then used to count the fragments mapping to each gene region. This generates a gene activity matrix where rows represent cells, columns represent genes, and each cell value indicates the number of fragments in the corresponding gene region. Finally, the gene activity scores are log-normalized to standardize the data.

- **Cicero** [13] calculates gene activity scores by assessing the chromatin accessibility at the promoter region and the regulatory potential of nearby chromatin peaks. It uses a correlation-based model to quantify the relationship between distal regulatory regions and the target gene's promoter. Specifically, Cicero identifies chromatin interactions by measuring the co-accessibility between the promoter and distal peaks, where high co-accessibility indicates potential gene activation.

- **Gene Scoring** [11] assigns each gene an accessibility score by aggregating the chromatin accessibility peaks around its transcription start site (TSS). The method applies an exponential decay function to weight peaks according to their distance from the TSS, with closer peaks given higher weights. This scoring system reflects the regulatory influence of proximal enhancers or other elements near the TSS on gene activity.

- **MAESTRO** [19] computes a gene activity score as a weighted sum of nearby cis-regulatory elements (REs), where the weight is determined by an exponentially decaying function of the distance between the regulatory elements (such as enhancers) and the target gene. This method quantifies the influence of nearby regulatory elements on gene activity, with closer elements having a stronger regulatory effect.

- **cisTopic** [1] applies Latent Dirichlet Allocation (LDA), a Bayesian topic modeling technique commonly applied in natural language processing, to identify cell states based on the distribution of topics across cells, and to explore cis-regulatory regions through the distribution of regions across topics. By combining the topic-cell distribution with the region-topic distribution, cisTopic calculates the probability of each region being active in a given cell (i.e., the predictive distribution). Finally, the gene activity score for each gene is derived by summing the probabilities of regions associated with that gene, typically those linked to known marker genes.

- **SnapATAC2** [14] Instead of fragment counts, SnapATAC2 uses counts of Tn5 insertion sites within gene bodies and normalizes these counts using a log-transformed count-per-million reads (CPM) approach. Additionally, SnapATAC2 employs a method based on Markov Affinity Graphs for data imputation and smoothing. This method takes into account the proximity of Tn5 insertion sites, weights the distance of these sites from the transcription start site (TSS), and also considers the influence of distal regulatory elements.

#### 1.3 Data preprocessing

To integrate scRNA-seq data with the gene activity score matrix from scATAC-seq data, both sharing the same gene features, we utilize a structured preprocessing pipeline implemented with the Scanpy package. The process starts by identifying HVGs using the *scanpy.pp.highly_variable_genes* function for each matrix individually. These HVGs from both datasets are then combined to form a unified set of HVGs for the integration. The subsequent normalization steps are performed separately for each dataset. Specifically, we use the *scanpy.pp.normalize_total* function for normalization, followed by log transformation with *scanpy.pp.log1p*, and scaling the data using *scanpy.pp.scale*. Finally, to reduce dimensionality, PCA embeddings are computed using the *scanpy.tl.pca* function, setting the number of principal components (npca) to 100 by default. This workflow ensures that both datasets are processed consistently, allowing for a more effective integration of the scRNA-seq and gene activity score data in subsequent analyses.

For integrating scRNA and single-cell protein modalities, the preprocessing steps for each modality are designed to handle their specific characteristics. For the protein data, we follow the standard procedures used in Seurat. The data is normalized using the NormalizeData() function, followed by scaling with the ScaleData() function. After these steps, PCA is applied to reduce dimensionality using the RunPCA() function. For the scRNA data, preprocessing can be performed using either the Scanpy or Seurat package. If using Seurat, the functions NormalizeData(), FindVariableFeatures(), ScaleData(), and RunPCA() are applied sequentially. Alternatively, using Scanpy, the preprocessing procedure is similar to that of scRNA and gene activity score matrix of scATAC data. Note that scRNA data is typically high-dimensional and sparse, which necessitates the selection of highly variable genes to reduce noise and improve the integration process. In contrast, protein data usually has fewer features (hundreds at most), which generally does not require feature selection, as its dimensionality is much lower.

#### 1.4 Additional experimental results

##### 1.4.1 Parameter settings

##### 1.4.2 Additional figures

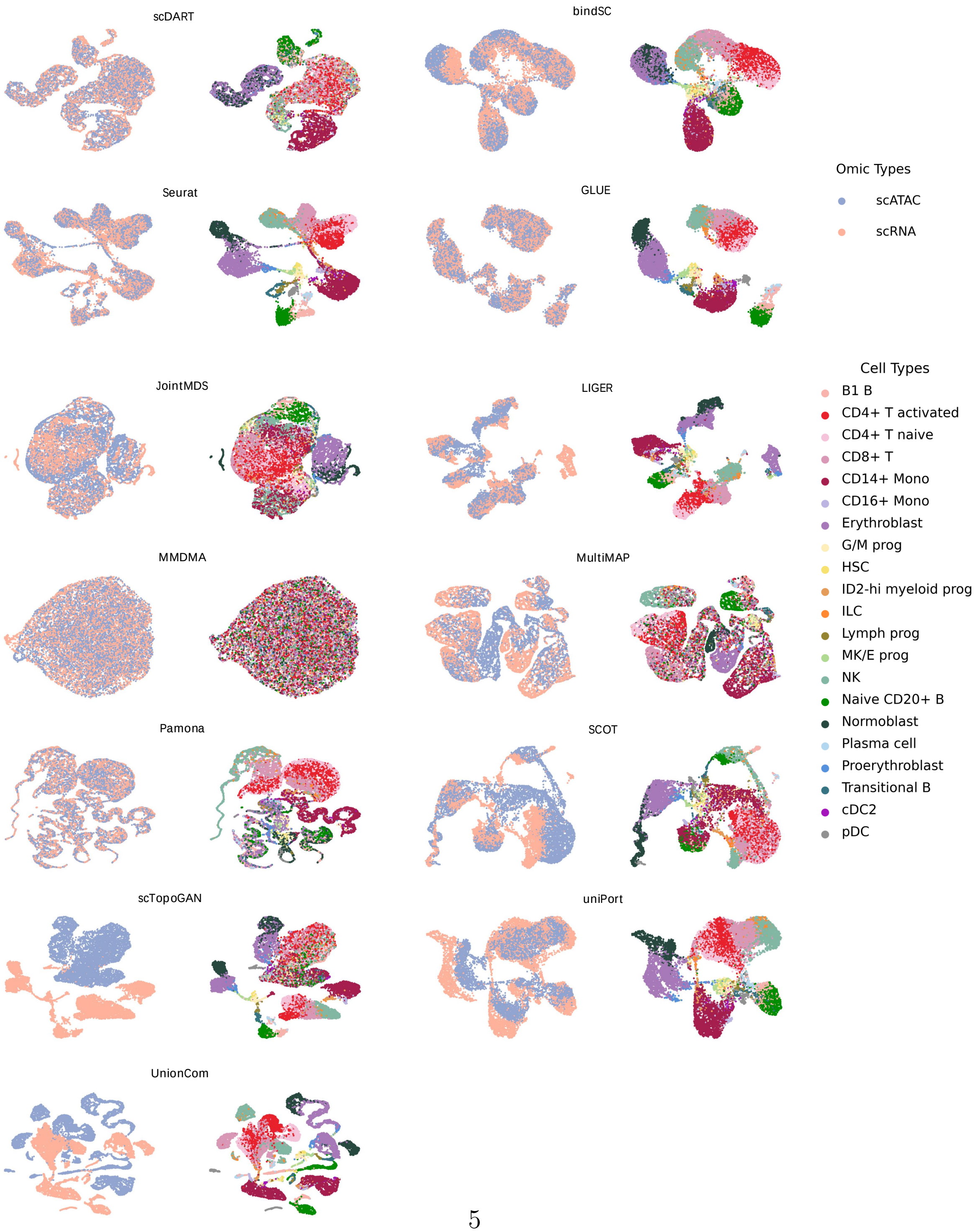




Fig. S1: UMAP visualizations of the integrated embeddings by different methods for BMMC data with cells colored based on omic types and cell types.

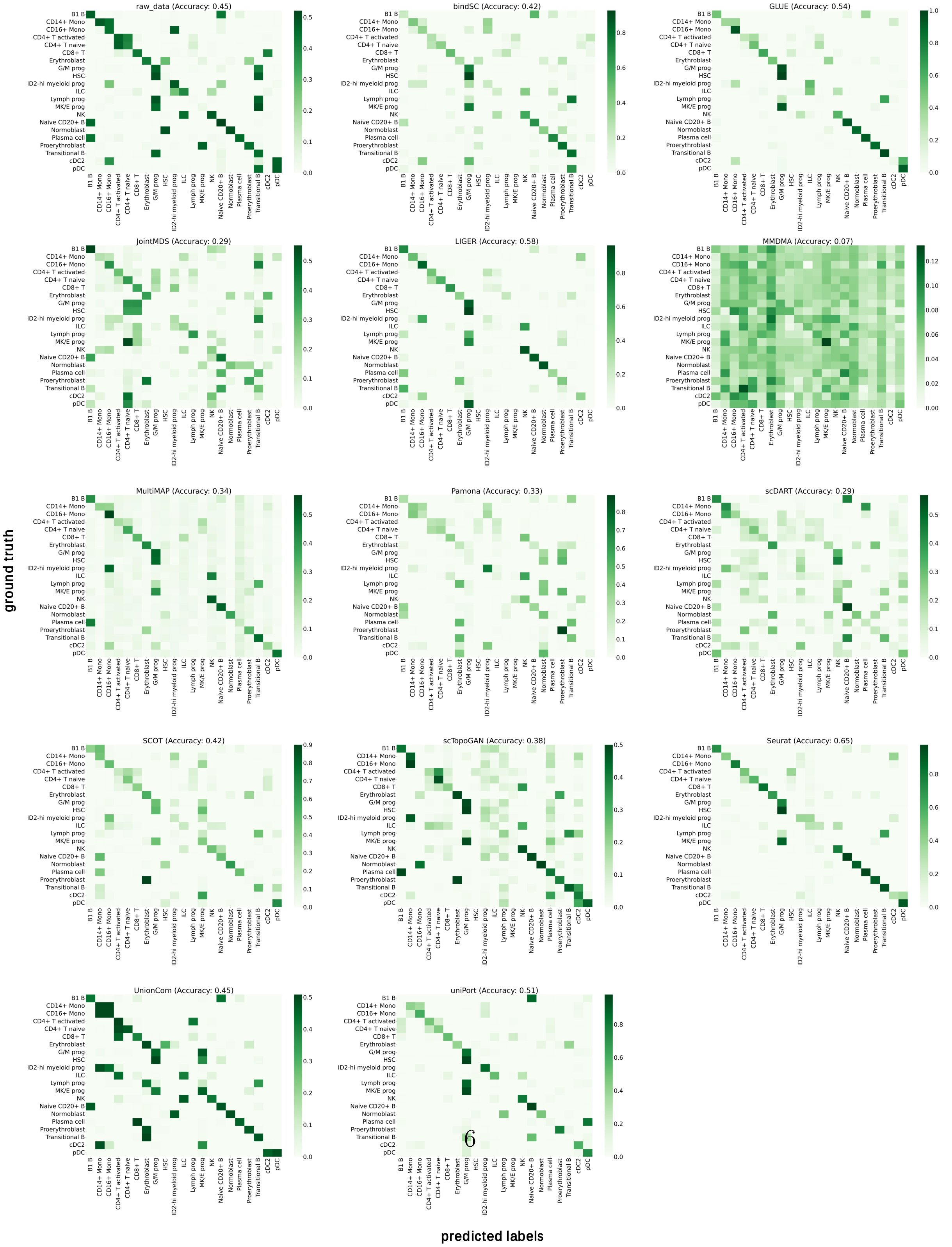


Fig. S2: Heatmap of the confusion matrix for different methods and the corresponding cell type prediction accuracy values in BMMC data, where rows represent the true cell types, columns represent the predicted cell types, and each element $i, j$ in the matrix represents the proportion of cells of type $i$ that are classified as type $j$.

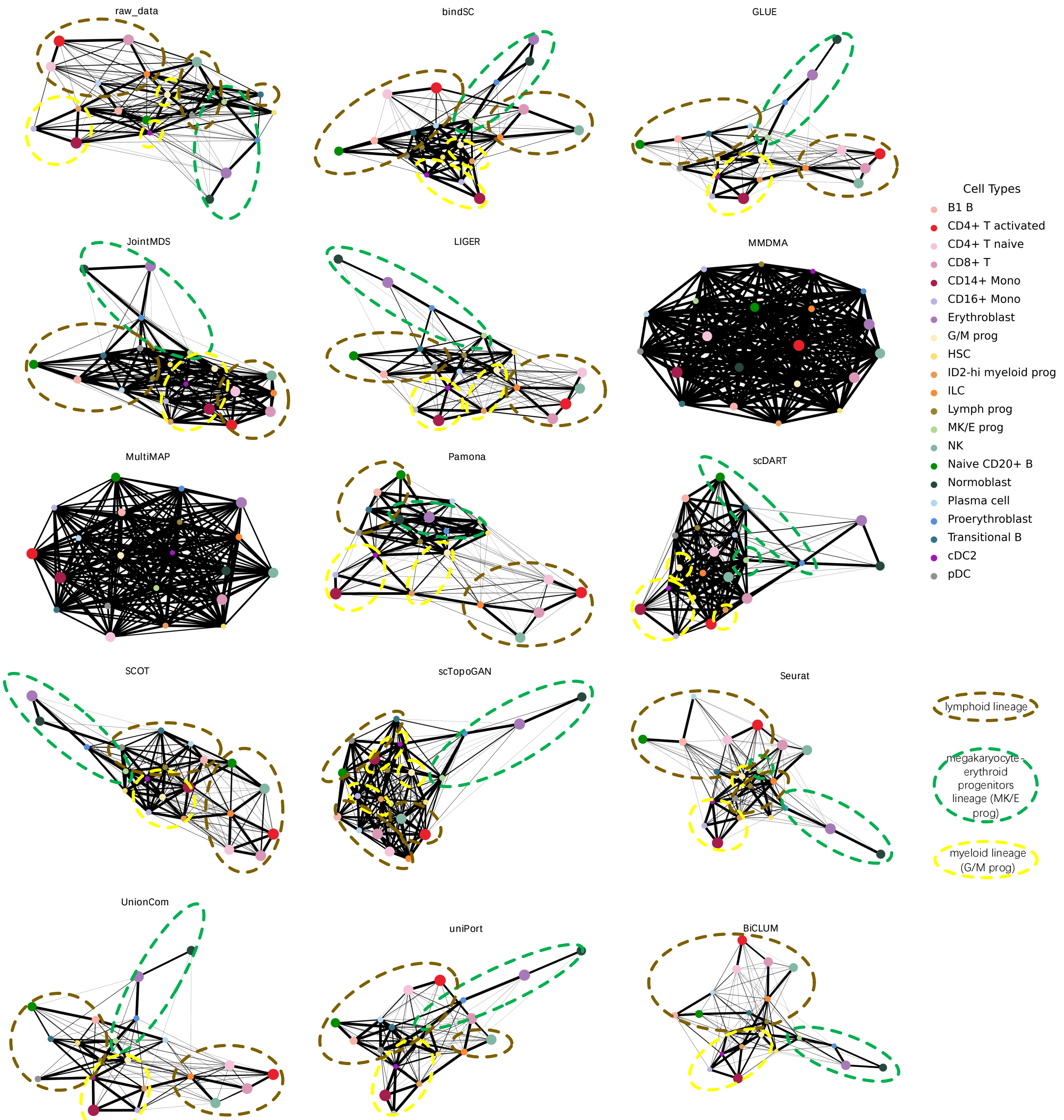


Fig. S3: PAGA trajectory visualizations for the BMMC data across different integration methods, where each node represents a cell type and the size is proportional to the number of cells in that type. Edges indicate potential lineage relationships, with the thickness representing the degree of connectivity between cell types.

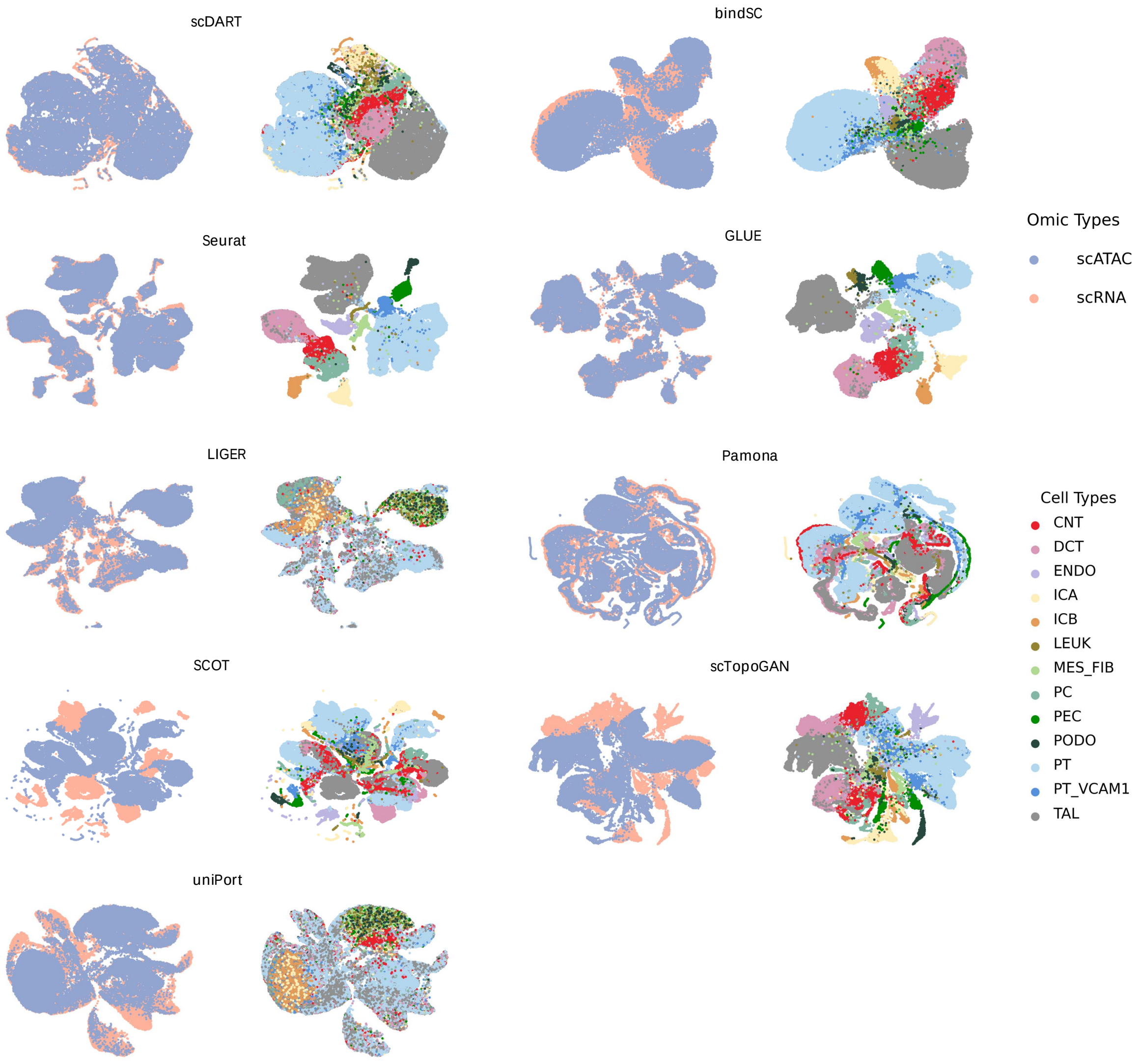


Fig. S4: UMAP visualizations of the integrated embeddings by different methods for Kidney data with cells colored based on omic types and cell types.

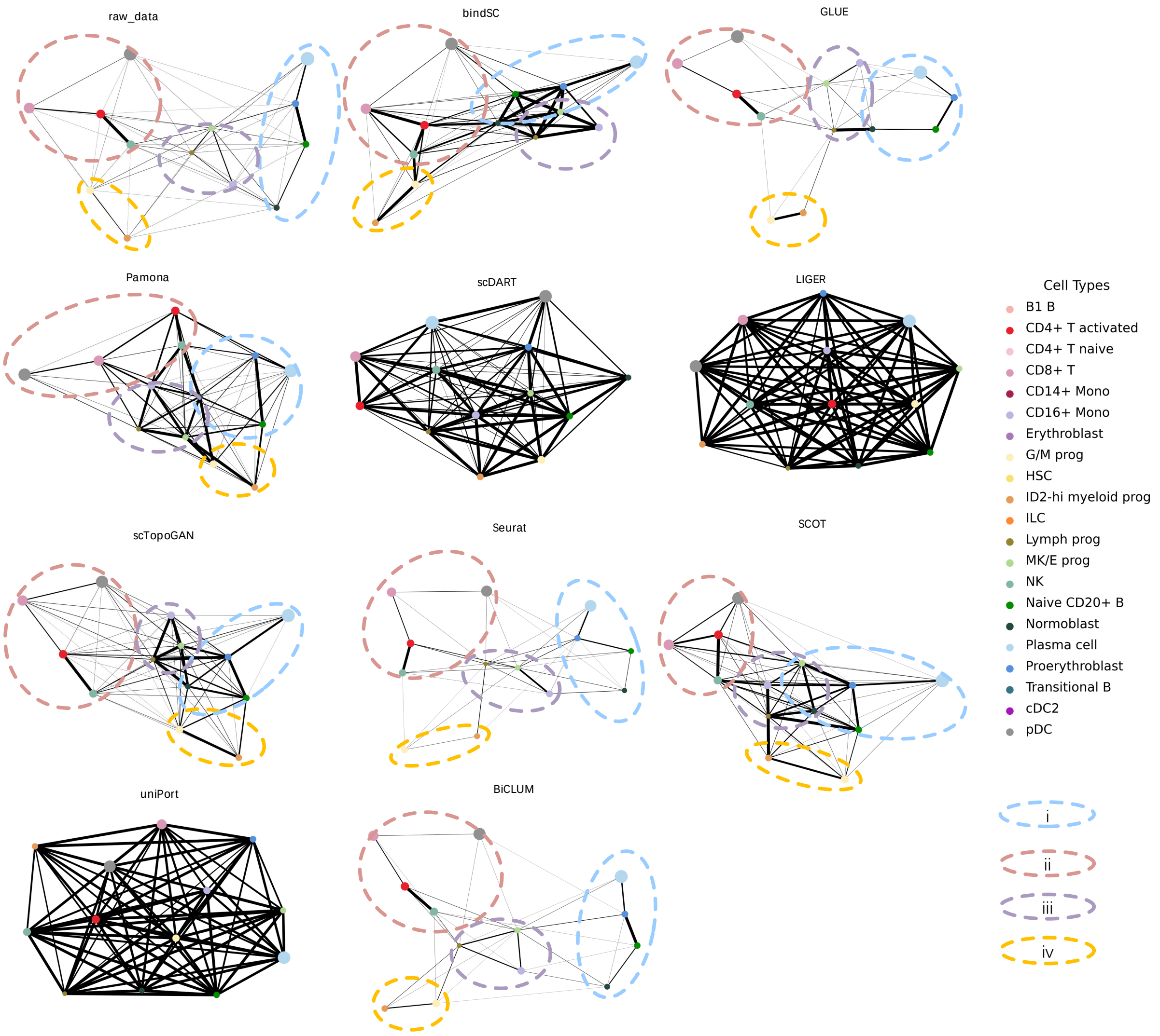


Fig. S5: PAGA trajectory visualizations for the Kidney data across different integration methods, where each node represents a cell type and the size is proportional to the number of cells in that type. Edges indicate potential lineage relationships, with the thickness representing the degree of connectivity between cell types.

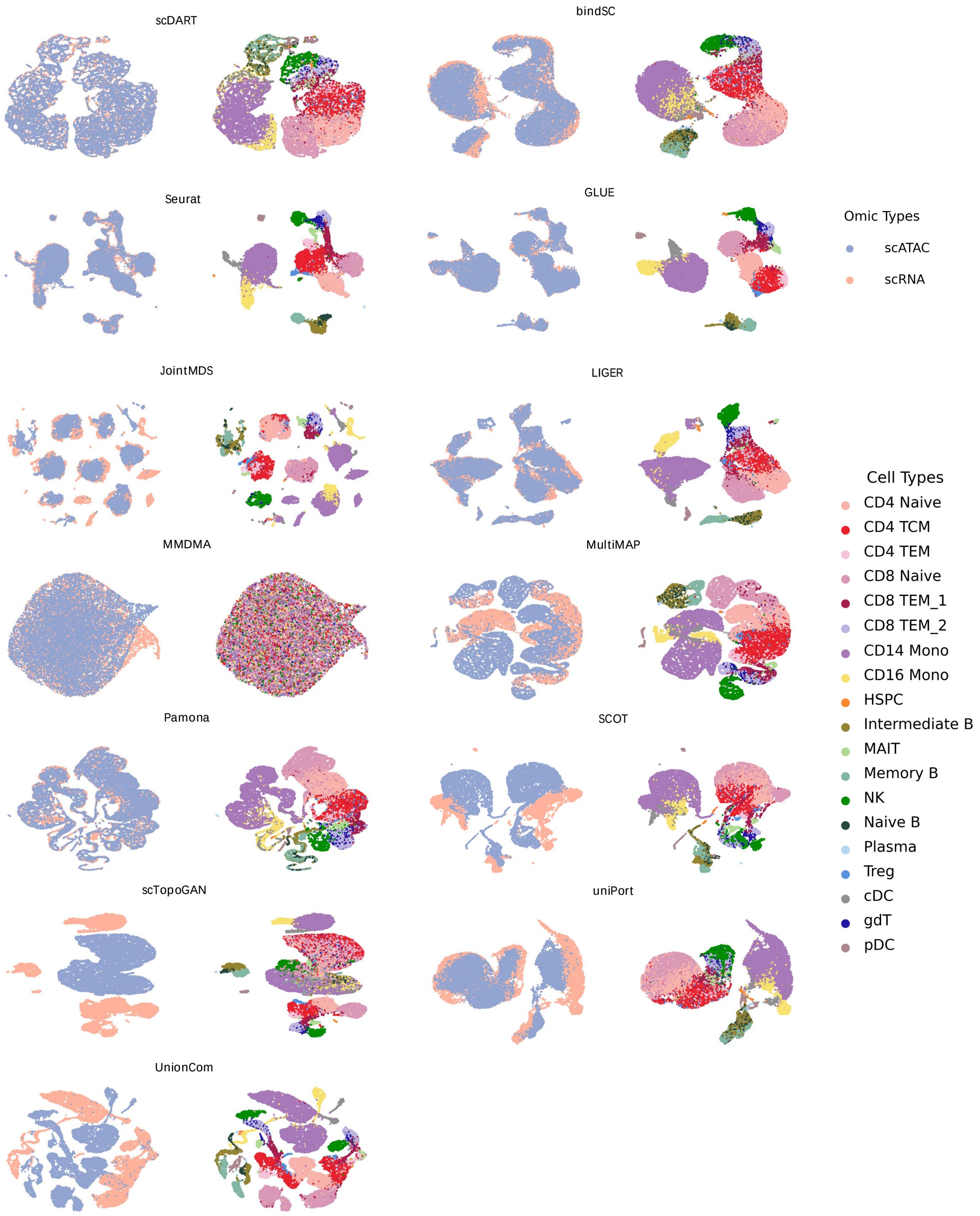




Fig. S6: UMAP visualizations of the integrated embeddings by different methods for PBMC (paired) data with cells colored based on omic types and cell types.

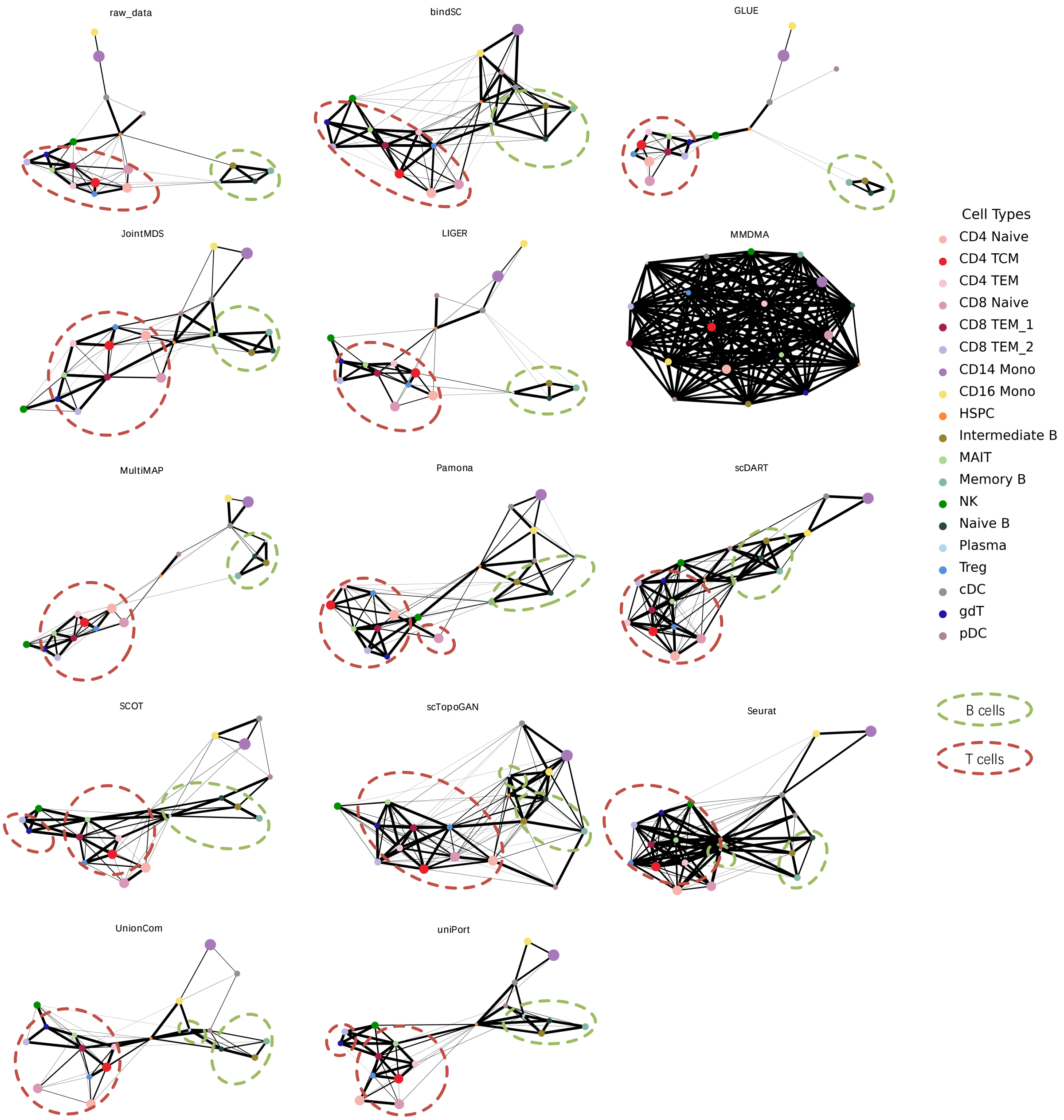


Fig. S7: PAGA trajectory visualizations for the PBMC (paired) data across different integration methods, where each node represents a cell type and the size is proportional to the number of cells in that type. Edges indicate potential lineage relationships, with the thickness representing the degree of connectivity between cell types.

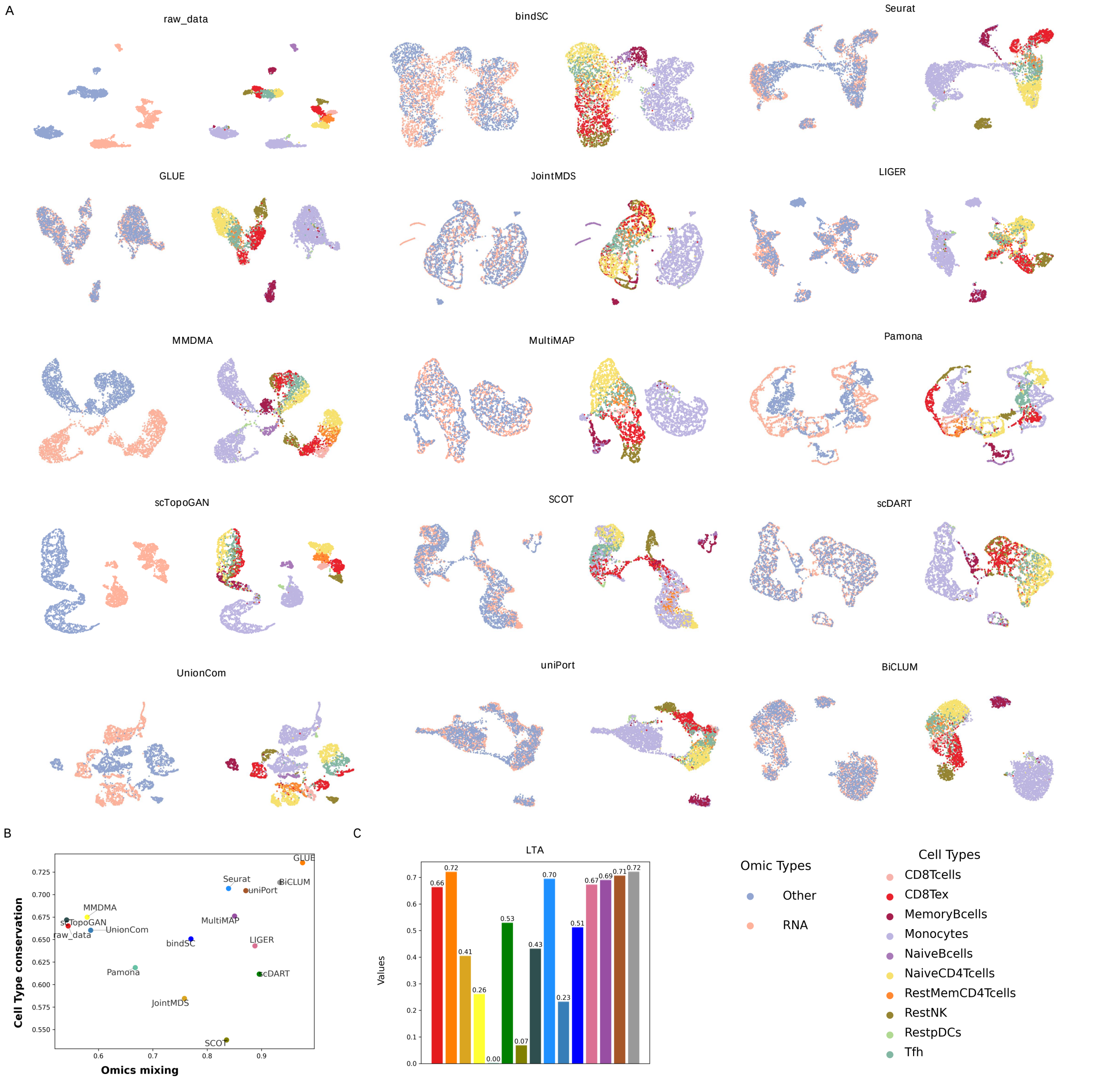


Fig. S8: Integrated results of PBMC (unpaired) data.(A) UMAP visualizations of the integrated embeddings by different methods for PBMC (unpaired) data with cells colored based on omic types and cell types. (B) Two evaluation metrics of omics mixing and biology conservation for multi-omics integration methods. (C) LTA across different integration methods.

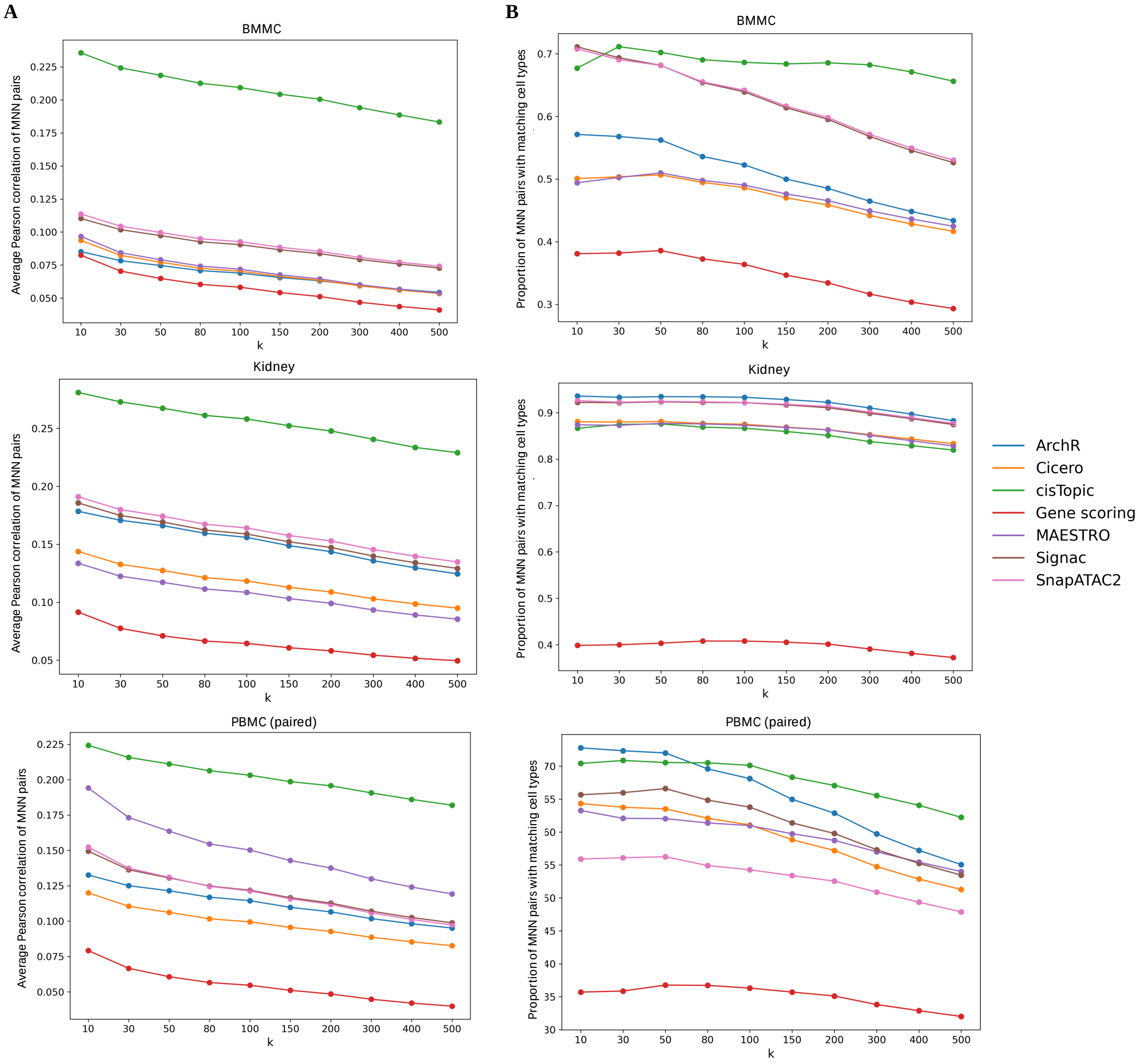


Fig. S9: Comparison of transformed matrices generated by different conversion methods for three datasets. MNN pairs were constructed based on these matrices and scRNA-seq data as $k$ increases. Evaluation was performed using (A) Average Pearson correlation of the constructed MNN pairs and (B) the proportion of MNN pairs with matching cell types.

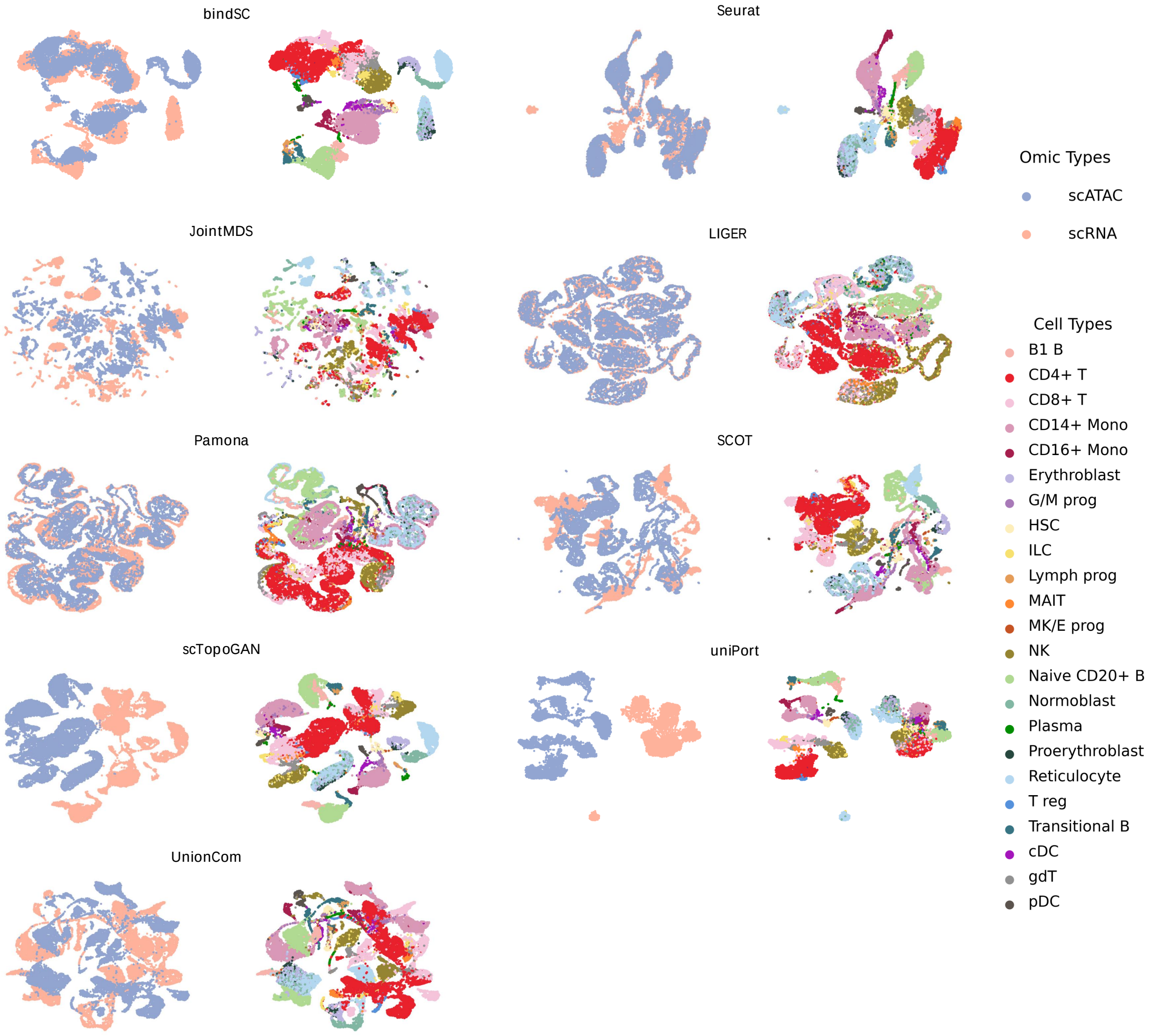


Fig. S10: UMAP visualizations of the integrated embeddings by different methods for BMCITE data for the two batches, s1d1 and s1d2, with cells colored based on omic types and cell types.

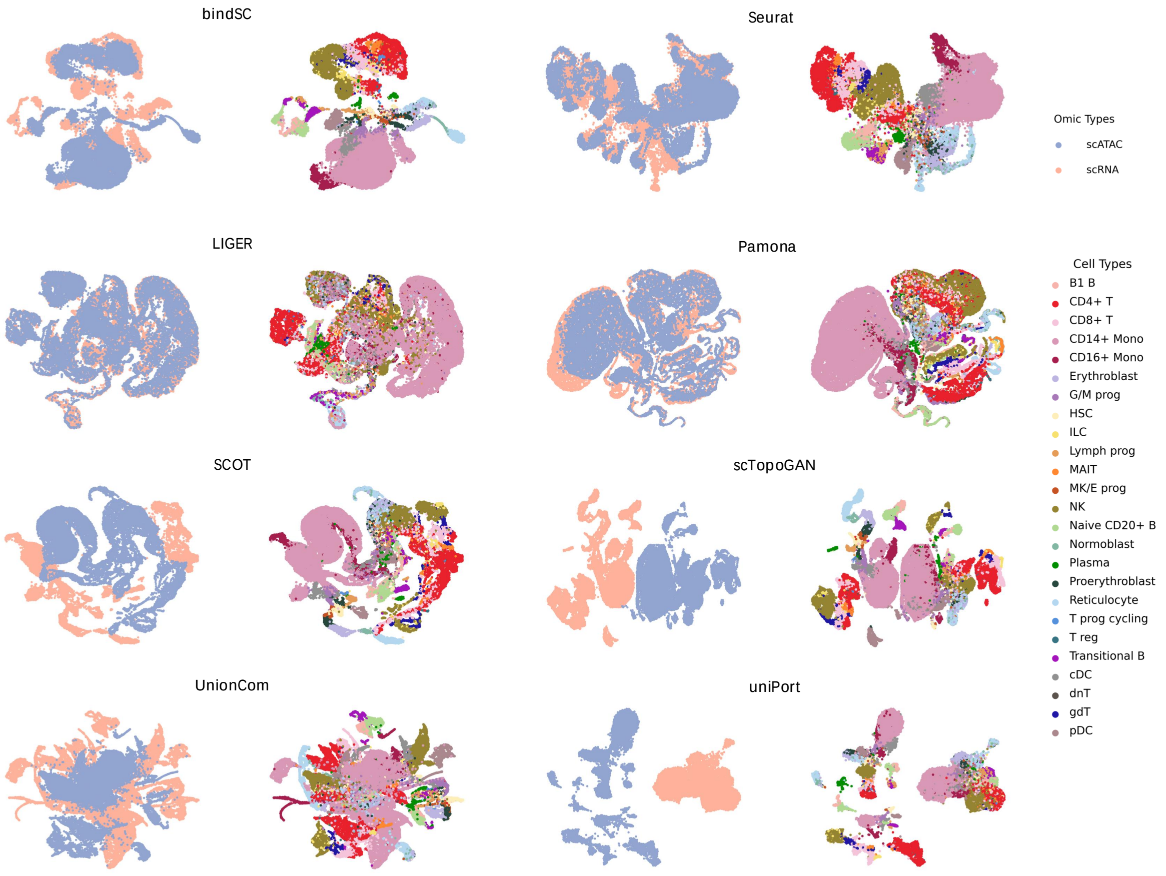


Fig. S11: UMAP visualizations of the integrated embeddings by different methods for BMCITE data for the two batches, s1d2 and s3d7, with cells colored based on omic types and cell types.

Table S1: The parameter settings for different datasets.

| data | method/batches | $\alpha$ | $\beta$ | $\tau_c$ | $\tau_f$ | $k_{mnn}$ | $d$ |
|---|---|---|---|---|---|---|---|
| BMMC | Signac | 1e4 | 1e4 | 0.5 | 0.5 | 200 | 50 |
| kidney | ArchR | 1e6 | 1e4 | 0.5 | 50 | 500 | 50 |
| PBMC (paired) | ArchR | 1e4 | 1e4 | 0.5 | 10 | 200 | 50 |
| PBMC (unpaired) | MAESTRO | 1e4 | 1e4 | 0.5 | 0.5 | 100 | 50 |
| BMCITE | s1d1_s1d2 | 1e3 | 0.1 | 100 | 50 | 500 | 25 |
| | s1d2_s3d7 | 1e4 | 1e6 | 100 | 50 | 500 | 25 |

[2] Kai Cao, Xiangqi Bai, Yiguang Hong, and Lin Wan. Unsupervised topological alignment for single-cell multi-omics integration. *Bioinformatics*, 36(Supplement_1):i48–i56, 2020.

[3] Kai Cao, Qiyu Gong, Yiguang Hong, and Lin Wan. A unified computational framework for single-cell data integration with optimal transport. *Nature Communications*, 13(1):7419, 2022.

[4] Kai Cao, Yiguang Hong, and Lin Wan. Manifold alignment for heterogeneous single-cell multi-omics data integration using pamona. *Bioinformatics*, 38(1):211–219, 2022.

[5] Zhi-Jie Cao and Ge Gao. Multi-omics single-cell data integration and regulatory inference with graph-linked embedding. *Nature Biotechnology*, 40(10):1458–1466, 2022.

[6] Dexiong Chen, Bowen Fan, Carlos Oliver, and Karsten Borgwardt. Unsupervised manifold alignment with joint multidimensional scaling. *arXiv preprint arXiv:2207.02968*, 2022.
